# Transcriptional reprogramming from innate immune functions to a pro-thrombotic signature upon SARS-CoV-2 sensing by monocytes in COVID-19

**DOI:** 10.1101/2022.04.03.486830

**Authors:** Allison K. Maher, Katie L. Burnham, Emma Jones, Laury Baillon, Claudia Selck, Nicolas Giang, Rafael Argüello, Charlotte-Eve Short, Rachael Quinlan, Wendy S. Barclay, Nichola Cooper, Graham P. Taylor, Emma E. Davenport, Margarita Dominguez-Villar

## Abstract

Alterations in the myeloid immune compartment have been observed in COVID-19, but the specific mechanisms underlying these impairments are not completely understood. Here we examined the functionality of classical CD14^+^ monocytes as a main myeloid cell component in well-defined cohorts of patients with mild and moderate COVID-19 during the acute phase of infection and compared them to that of healthy individuals. We found that *ex vivo* isolated CD14^+^ monocytes from mild and moderate COVID-19 patients display specific patterns of costimulatory and inhibitory receptors that clearly distinguish them from healthy monocytes, as well as altered expression of histone marks and a dysfunctional metabolic profile. Decreased NFκB activation in COVID-19 monocytes *ex vivo* is accompanied by an intact type I IFN antiviral response. Subsequent pathogen sensing *ex vivo* led to a state of functional unresponsiveness characterized by a defect in pro-inflammatory cytokine expression, NFκB-driven cytokine responses and defective type I IFN response in moderate COVID-19 monocytes. Transcriptionally, COVID-19 monocytes switched their gene expression signature from canonical innate immune functions to a pro-thrombotic phenotype characterized by increased expression of pathways involved in hemostasis and immunothrombosis. In response to SARS-CoV-2 or other viral or bacterial components, monocytes displayed defects in the epigenetic remodelling and metabolic reprogramming that usually occurs upon pathogen sensing in innate immune cells. These results provide a potential mechanism by which innate immune dysfunction in COVID-19 may contribute to disease pathology.

## Main text

COVID-19 is a respiratory tract infection caused by severe acute respiratory syndrome corona virus 2 (SARS-CoV-2). In unvaccinated individuals, the majority of infections are mild or asymptomatic, but 15% of patients develop moderate to severe disease requiring hospitalisation, and 5% develop critical disease with life-threatening pneumonia, acute respiratory distress syndrome (ARDs) and septic shock^1^. During the acute phase of infection, myeloid cells including monocytes and macrophages are the most enriched immune cell types in the lungs of COVID-19 patients and play a major role in the pathogenicity of the disease^2,3^. Moreover, contrasting observations regarding the development of cytokine storms vs. immunosuppression^4,5^ and the overactive or deficient type I IFN response in the lungs and in peripheral blood^6-11^ have been described for the role of myeloid cells in COVID-19^12^. Despite these apparent contrasting works, most studies have observed dysregulated innate immune responses and reduced expression of human leukocyte antigen DR isotype (HLA-DR) by circulating myeloid cells, which is considered a marker of immune suppression^10,13-17^.

Monocytes are blood-circulating, phagocytic, innate immune leukocytes with important functions in pathogen sensing, and innate and adaptive immune response activation during viral infection^18^. Despite their heterogeneity^19^, monocytes are broadly classified into three subsets based on the expression of CD14 and CD16 into classical (CD14^+^CD16^-^), intermediate (CD14^+^CD16^+^), and nonclassical (CD14^low^CD16^+^) monocytes^18^. During viral infection, circulating monocytes infiltrate affected tissues and differentiate into inflammatory macrophages and dendritic cells (DCs)^20^, contributing to pathogen clearance and tissue regeneration.

Here we deeply examined the phenotype and functionality of the main monocyte population in humans, i.e. classical CD14^+^ monocytes, in patients with COVID-19 and compared them to those of healthy individuals. We found that *ex vivo* isolated CD14^+^ monocytes from mild and moderate COVID-19 patients are phenotypically different from monocytes from healthy individuals, displaying differential expression of costimulatory receptors and MHC molecules, epigenetic alterations and a dysfunctional metabolic profile that is accompanied by decreased *ex vivo* NFκB activation, while maintaining an intact type I IFN antiviral response. Subsequent pathogen sensing *ex vivo* led to a state of functional unresponsiveness that correlated transcriptionally with that of a endotoxin-induced tolerance signature. Moreover, monocytes switched their gene expression signature from canonical innate immune functions to a pro-thrombotic phenotype characterized by increased expression of pathways involved in immunothrombosis. In response to SARS-CoV-2 or other viral or bacterial components, monocytes displayed decreased expression of type I IFN responses, decreased pro-inflammatory cytokine production and costimulatory receptor expression and defects in the epigenetic remodelling and metabolic reprogramming that usually occurs upon pathogen sensing. These results provide a potential mechanism by which innate immune dysfunction in COVID-19 contributes to disease progression and identifies potential therapeutic targets.

### Phenotypic and epigenetic alterations in COVID-19 monocytes

Global alterations in innate immune cell phenotypes have been identified in severe COVID-19^11,21-23^. As the main human monocyte population, we focused on deeply characterizing the *ex vivo* phenotype of classical CD14^+^ monocytes in uninfected healthy individuals and patients with COVID-19 presenting with mild or moderate symptoms (1-2 or 3-4 WHO ordinal scale for COVID-19 severity, respectively) during the acute phase of disease. The battery of markers examined by high dimensional flow cytometry included MHC molecules and costimulatory and coinhibitory receptors (Figure 1). Dimensionality reduction tools demonstrated that while some overlap in the global phenotypes was observed among the three study groups, monocytes from healthy individuals were clearly distinct from both mild and moderate COVID-19 on a tSNE plot (Figure 1a). In addition, COVID-19 monocytes could also be distinguished based on disease severity, with main cell clusters for both disease severity groups mapping separately on the tSNE plots. Moderate COVID-19 monocytes expressed decreased levels of HLA-DR, in agreement with previous reports^10,17^, but in contrast, they displayed increased expression of HLA-ABC compared to both mild disease and uninfected individuals, suggesting a skewed trend towards class I antigen presentation (Figure 1b). In addition, moderate COVID-19 monocytes expressed increased levels of the c-type lectin CD301. The decreased expression of the costimulatory receptor CD86 and increased expression of the inhibitory receptors TIM-3^24^ and PD-1^25^ on moderate COVID-19 monocytes suggest an altered activation profile skewed towards an inhibitory phenotype. Furthermore, there were significant differences in the expression of certain markers on mild vs. moderate COVID-19 monocytes. For example, downregulation of HLA-DR and CD86 and upregulation of TIM-3 and HLA-ABC compared to healthy monocytes were only significant in moderate but not on mild COVID-19 monocytes, and the increased expression of CD80 in mild COVID-19 compared to healthy monocytes was not apparent in moderate COVID-19. These results suggest a more profound dysfunction in moderate than in mild COVID-19 monocytes.

**Figure 1.**
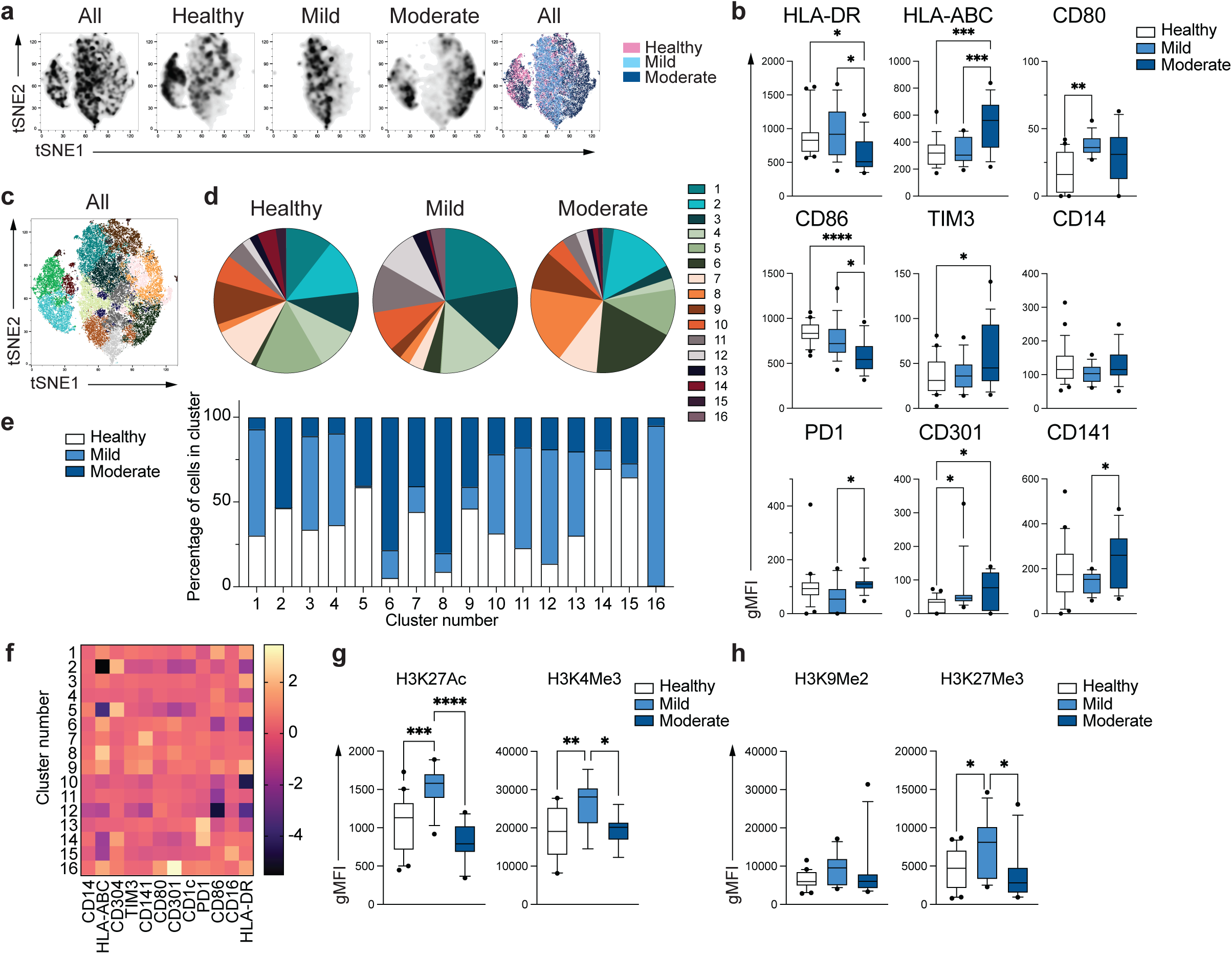
Unique phenotype of COVID-19 monocytes. **a**. tSNE plots obtained from a concatenated sample consisting of PBMC from n=15 healthy individuals, n=15 mild and n=15 moderate COVID-19 patients. **b**. Box and whiskers plots summarizing the median gMFI of the receptors analyzed. The box extends from the 25^th^ to the 75^th^ percentile and the whiskers are drawn down to the 10^th^ percentile and up to the 90^th^ percentile. Points below and above the whiskers are drawn as individual points (n=25 healthy, n=15 mild and n=17 moderate COVID-19 individuals). **c**. tSNE plots depicting the cell clusters identified by Phenograph from the concatenated sample in **a. d**. Pie charts show the fraction of cells within each identified cell cluster in each patient group. **e**. Bars graph show the distribution (percentage) of cells from each patient group in each identified cell cluster. **f**. Heatmap of the expression of receptors per cell cluster displayed as modified z-scores using median values. **g** and **h**. Summary of expression of activating (**g**) and repressive (**h**) histone marks in monocytes from healthy individuals (n=20), mild (n=15) and moderate (n=11) COVID-19 patients. One-way ANOVA with Tukey’s correction for multiple comparisons for **b, g, h**. *P<0.05, **p<0.005, ***p<0.001, ****p<0.0001.

To further define and quantify the phenotypic differences observed between healthy individuals and COVID-19 patients, we applied clustering algorithms using the 12 phenotypic markers previously examined. Cell clustering identified 16 different subpopulations of monocytes that were distinctively distributed in healthy and COVID-19 monocytes (Figure 1c, d), with 11 clusters containing more than 88% of the total cells analyzed (Supplementary Figure 1). Interestingly, expansion of specific monocyte subpopulations were different in mild and moderate COVID-19 monocytes, and while mild monocytes, in contrast to healthy monocytes, predominantly contained clusters 1, 3 and 4 and did not contain clusters 2 and 5, monocytes from moderate COVID-19 patients significantly had reduced frequency of cells from clusters 1, 3 and 4, and contained expanded clusters 6 and 8 (Figure 1d and Supplementary Table 2). As a consequence, the distribution of cells from healthy, mild and moderate COVID-19 monocytes was clearly different in each cluster, and while some cell clusters were composed of cells from all disease groups, such as clusters 10, 11 and 13, other clusters predominantly contained cells from one or two particular disease groups. For example, clusters 1, 3, 4, 12 and 16 were predominantly composed of cells from mild patients, while clusters 6 and 8 predominantly contained moderate COVID-19 monocytes and were almost absent in monocytes from healthy individuals (Figure 1e). Normalized expression levels of the markers defining each cluster demonstrated that the phenotype of cluster 6 was mostly driven by downregulation of CD86 and HLA-DR, while that of cluster 8 was mostly driven by the increased expression of HLA-ABC (Figure 1f). Collectively, these results reveal that distinct populations of circulating monocytes are enriched in mild and moderate COVID-19 patients.

As a measurement of global differences in the patterns of activation/repression of gene expression we looked at the protein expression of histone marks associated with active gene transcription (H3K27Ac and H3K4Me3^26,27^, Figure 1g) and gene repression (H3K9Me2 and H3K27Me3^26,27^, Figure 1h) in monocytes from healthy individuals and patients with COVID-19 *ex vivo*. Significant differences in the expression of epigenetic marks associated with activation of gene expression were found. Monocytes from mild COVID-19 patients displayed increased levels of both H3K27Ac and H3K4Me3 compared to healthy individuals as expected considering the *in vivo* pathogen sensing and subsequent activation of innate immunity by an ongoing viral infection^28^. However, moderate COVID-19 monocytes failed to increase H3K27Ac and H3K4Me3 expression and displayed similar levels to those of healthy individuals (Figure 1g). Moreover, while no differences were observed in the expression of the repressive mark H3K9Me2, the increased H3K27Me3 observed in mild COVID-19 monocytes was not observed in moderate COVID-19. These results suggest that the epigenetic remodeling associated with virus sensing and subsequent activation of innate immunity is defective in moderate COVID-19 monocytes.

### *Ex vivo* RNA-seq uncovers metabolic dysfunction in moderate COVID-19 monocytes

The fundamental differences in the phenotype and epigenetic marks in moderate COVID-19 monocytes compared to those of healthy individuals led us to investigate in depth the gene expression profile of *ex vivo* isolated classical CD14^+^ monocytes from patients with moderate COVID-19 and compare them with those of healthy individuals (Figure 2). Principal component analysis (PCA) applied to examine the global distribution of gene expression profiles from COVID-19 monocytes (n=10) and healthy individuals (n=6) demonstrated a clear separation between groups along PC1 (Figure 2a), with genes encoding a number of soluble factors, chemokines and class II molecules as the main genes contributing to the separation between healthy and COVID-19 monocytes (Supplementary Figure 2). Differential gene expression analysis yielded 422 upregulated and 187 downregulated genes (≥ 1.5-fold change, FDR<0.05) in COVID-19 monocytes compared to healthy controls (Figure 2b). We used these genes to perform a pathway enrichment analysis with XGR^29^ and pathway annotations from Reactome to gain insight on potential pathways differentially expressed in COVID-19 monocytes (Supplementary Figure 3). Interestingly, pathway enrichment identified glycolysis as the most enriched pathway in COVID-19 monocytes together with metabolism of lipids and lipoproteins. Moreover, the presence of interferon signaling and cytokine signaling in the list of enriched pathways was in agreement with previous reports on the role of these two pathways in COVID-19 pathogenesis^6,17,23^ (Supplementary Figure 3 and Supplementary Table 3).

**Figure 2.**
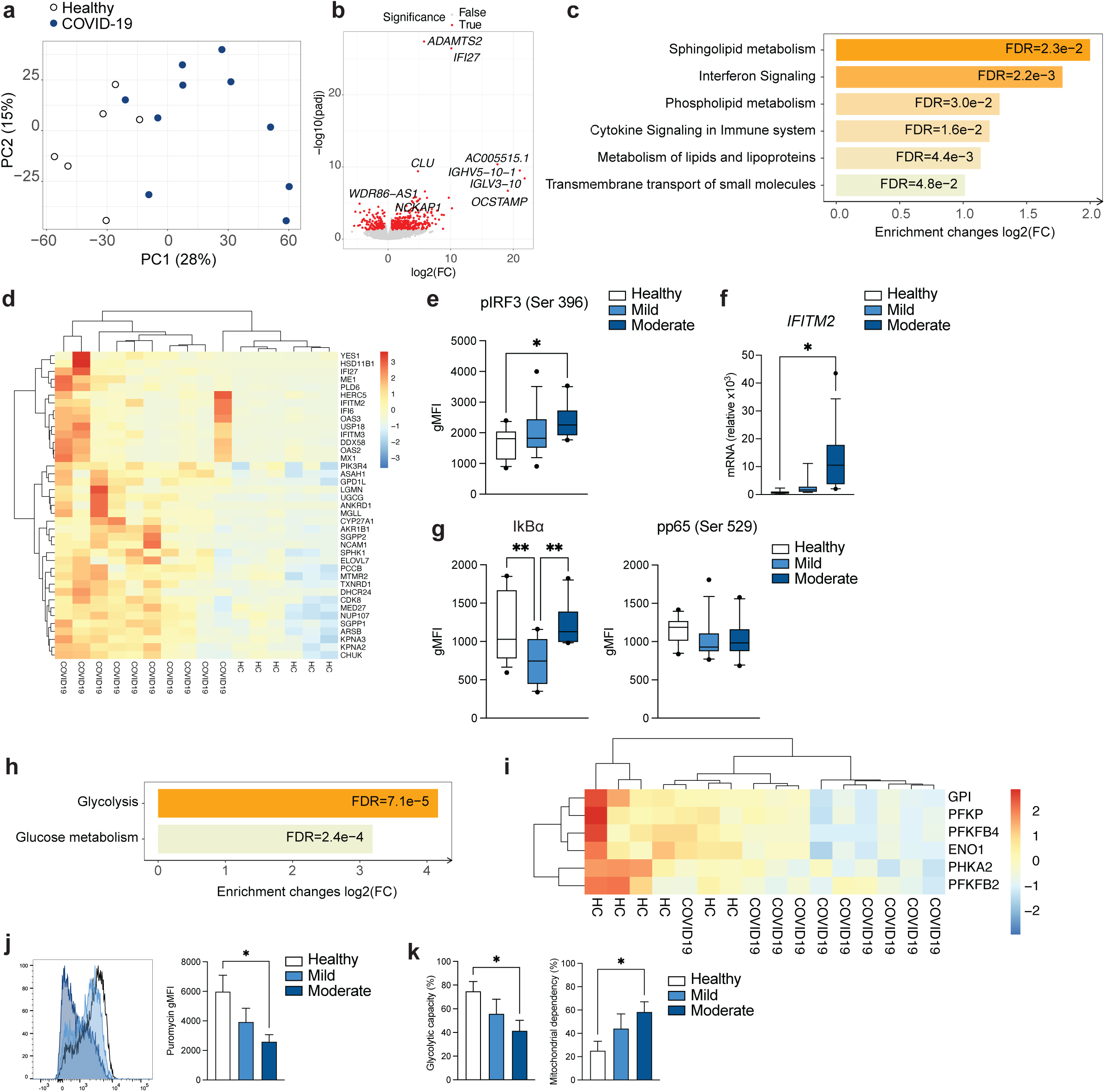
Gene expression signature of COVID-19 monocytes *ex vivo*. **a**. Principal component analysis (PCA) of the gene expression data computed from all genes from *ex vivo* healthy individual (white dots) and moderate COVID-19 (blue dots) monocyte samples. PC2 plotted against PC1 to explore overall variation across samples. The variance explained by each component is stated in brackets. **b**. Volcano plot of differentially expressed genes for ex vivo COVID-19 vs healthy monocytes. Red coloring shows genes with fold change ≥1.5 and FDR<0.05. **c**. Bar plots depict significantly enriched (FDR<0.05) pathways from Reactome for COVID-19 vs. healthy individual monocytes using upregulated genes in COVID-19 vs healthy (≥1.5 fold increase, FDR<0.05), with the fold enrichment plotted on the x axis as log_2_ (FC) and the bars labelled with the adjusted p value. **d**. Significantly upregulated genes in the COVID-19 vs healthy monocyte contrast that are members of the pathways in **c**, shown in a heatmap. Gene expression values are scaled by row, with red indicating relatively high expression and blue low expression. Both rows and columns are clustered using Euclidean distance and Ward’s method. **e**. Phospho-IRF3 (Ser 396) expression measured by flow cytometry and plotted as gMFI for healthy (n=14), mild (n=15) and moderate (n=10) COVID19 monocytes. **f**. *IFITM2* relative gene expression (to *GAPDH*) measured by real-time PCR in sorted CD14^+^ monocytes from healthy individuals (n=7), mild (n=7) and moderate (n=13) COVID-19. **g**. Iκ Bα (left) and phospho-NFκB p65 (right) expression measured by flow cytometry as gMFI in healthy individuals (n=14), mild (n=15) and moderate (n=10) COVID-19 monocytes. **h**. Bar plots depict significantly enriched (FDR<0.05) pathways from Reactome for COVID-19 vs. healthy individual monocytes, using downregulated genes in COVID-19 vs. healthy (≥1.5 fold decrease, FDR<0.05), with the fold enrichment plotted on the x axis as log_2_ (FC) and the bars labelled with the adjusted p value. **i**. Significantly downregulated genes in the COVID-19 vs. healthy monocyte contrast that are members of the pathways in **h**, shown in a heatmap. Gene expression values are scaled by row, with red indicating relatively high expression and blue low expression. Both rows and columns are clustered using Euclidean distance and Ward’s method. **j**. Representative example of *ex vivo* expression of puromycin in CD14^+^ monocytes measured by flow cytometry (left) and summary of puromycin gMFI on healthy individuals (n=10), mild (n=8) and moderate (n=10) COVID-19 monocytes (right). **k**. Glycolytic capacity (left) and mitochondrial dependency (right) of monocytes from healthy individuals (n=10), mild (n=8) and moderate (n=10) COVID-19 monocytes *ex vivo*. One-way ANOVA with Tukey’s test for multiple comparisons in **e, f, g, j, k**. *p<0.05, **p<0.005.

We subsequently examined the directionality of expression of the enriched pathways by analyzing downregulated genes and upregulated genes separately. Pathway enrichment analysis of genes significantly upregulated (≥ 1.5-fold change, FDR<0.05) in COVID-19 compared to healthy individuals demonstrated a significant increase in the metabolism of a number of lipids, including sphingolipids, phospholipids and lipoproteins. Other upregulated pathways in COVID-19 monocytes included interferon signaling, cytokine signaling and transmembrane transport of small molecules. Heatmap showing the top 40 upregulated genes from the enriched pathways demonstrated a somewhat variable expression patterns among COVID-19 monocytes and included a number of type I interferon-stimulated genes (*IFI27, IFITM2, IFI6, IFITM3, MX1*), metabolic enzymes (*ASAH1, CYP27A1, SGPP2, SPHK1*) and others (Figure 2d). Interestingly, the highest expressed IFN-related gene was *IFI27*, which has been suggested as a biomarker of early SARS-CoV-2 infection^30^. The increased type I IFN gene signature in COVID-19 monocytes was confirmed by the increased *ex vivo* phospho-IRF3 protein expression in moderate COVID-19 patients compared to healthy individuals (Figure 2e) and by the increased expression of *IFITM2* as an IFN-stimulated gene, measured by real-time PCR in an expanded cohort of mild and moderate COVID-19 patients (Figure 2f). NFκB activation was examined *ex vivo* indirectly by Iκ Bα expression and directly by phosphorylation of the p65 NFκB subunit, as a readout for cytokine signaling^31,32^. While mild, unlike moderate COVID-19 monocytes displayed a decrease in the expression of IκBα compared to that of healthy individual monocytes, neither mild or moderate COVID-19 monocytes displayed an increased expression of phospho-p65 NFκB, suggesting that other additional mechanisms may be regulating the activation of NFκB, and that NFκB-driven cytokine responses may be altered in patients with COVID-19, in agreement with the lack of increased pro-inflammatory cytokine expression by COVID-19 monocytes (Figure 2c) and with previous single cell transcriptomic data of acute COVID-19 PBMC^33^. Moreover, several of the genes contributing to the “Cytokine signaling” pathway enrichment (Figure 2c) were interferon-stimulated genes (Supplementary Table 4).

We subsequently selected the set of significantly downregulated genes (≥1.5 fold decrease, FDR<0.05) in COVID-19 monocytes to perform pathway enrichment. The only pathway that was significantly downregulated in COVID-19 monocytes was glycolysis (Figure 2h, I and Supplementary Table 5). This metabolic profile with increased metabolism of lipids (Figure 2c) and decreased glycolysis was unexpected, as glycolysis is an important driver of innate immune cell function during the recognition of pathogens^34^. We used SCENITH^TM35^ to metabolically profile CD14^+^ monocytes from COVID-19 patients and healthy controls *ex vivo*. SCENITH™ uses protein synthesis as a measurement of global metabolic activity. Puromycin incorporation is used as a reliable readout of protein synthesis levels (and therefore metabolic activity) *in vitro* and *in vivo*. In agreement with the pathway enrichment results, *ex vivo* puromycin incorporation was significantly decreased in moderate COVID-19 monocytes (Figure 2j) compared to healthy individuals, suggesting decreased metabolic activity. Moreover, the glycolytic capacity of COVID-19 monocytes was significantly decreased in moderate patients and correlated with disease severity (Figure 2k), and this was accompanied by a concomitant increase in metabolic dependency in monocytes from moderate COVID-19 patients. The decreased metabolic activity and glycolytic capacity was further confirmed by Seahorse analysis of extracellular acidification rate and oxygen consumption rate as readouts for glycolysis and oxidative phosphorylation, respectively (Supplementary Figure 4).

These data suggest that monocytes from COVID-19 patients with moderate disease display epigenetic alterations and a dysfunctional metabolic profile that is accompanied by decreased NFκB activation, while maintaining intact type I IFN antiviral responses.

### COVID-19 monocytes display impaired pathogen sensing and activation mechanisms *ex vivo*

The dysfunctional metabolic profile with a downregulation of glycolysis and the defective activation of NFκB, both pathways heavily involved in the activation of innate immune cells upon virus encounter^32,34^, led us to examine the functional capacity of monocytes to sense and respond to SARS-CoV2 *ex vivo* (Figure 3). Stimulation of CD14^+^ monocytes from healthy individuals with SARS-CoV-2 led to a significant increase in both TNF and IL-10 production (Figure 3a). However, COVID-19 monocytes significantly produced less TNF as compared to healthy monocytes, while no differences were observed in IL-10 expression (Figure 3b). Moreover, the defect in TNF production upon stimulation was not SARS-CoV-2-specific, as stimulation with common cold coronaviruses or bacterial lipopolysaccharide (LPS) also led to significantly reduced TNF production compared to monocytes from healthy individuals (Figure 3c). In addition, the expression of CD40 (Figure 3d), which is important for monocyte effector function and is upregulated after virus sensing^36^, was increased in monocytes from healthy individuals but not on COVID-19 monocytes (Figure 3e). This decreased expression was confirmed after stimulation with common cold coronaviruses or LPS (Figure 3f), suggesting that the activation defects in COVID-19 monocytes in response to pathogen sensing were not specific to SARS-CoV-2. In addition to CD40, we also examined the expression of other cell surface receptors involved in antigen presentation and activation of T cells. (Figure 3g) HLA-DR expression levels were not further upregulated upon SARS-CoV-2 stimulation in any of the patient groups, and stimulation still maintained the differences in expression observed *ex vivo* among groups (Figure 1b). Moreover, while CD80 was significantly upregulated in healthy, mild and moderate COVID-19 monocytes after SARS-CoV-2 stimulation, only healthy monocytes increased the expression of CD86 after stimulation (Figure 3g).

**Figure 3.**
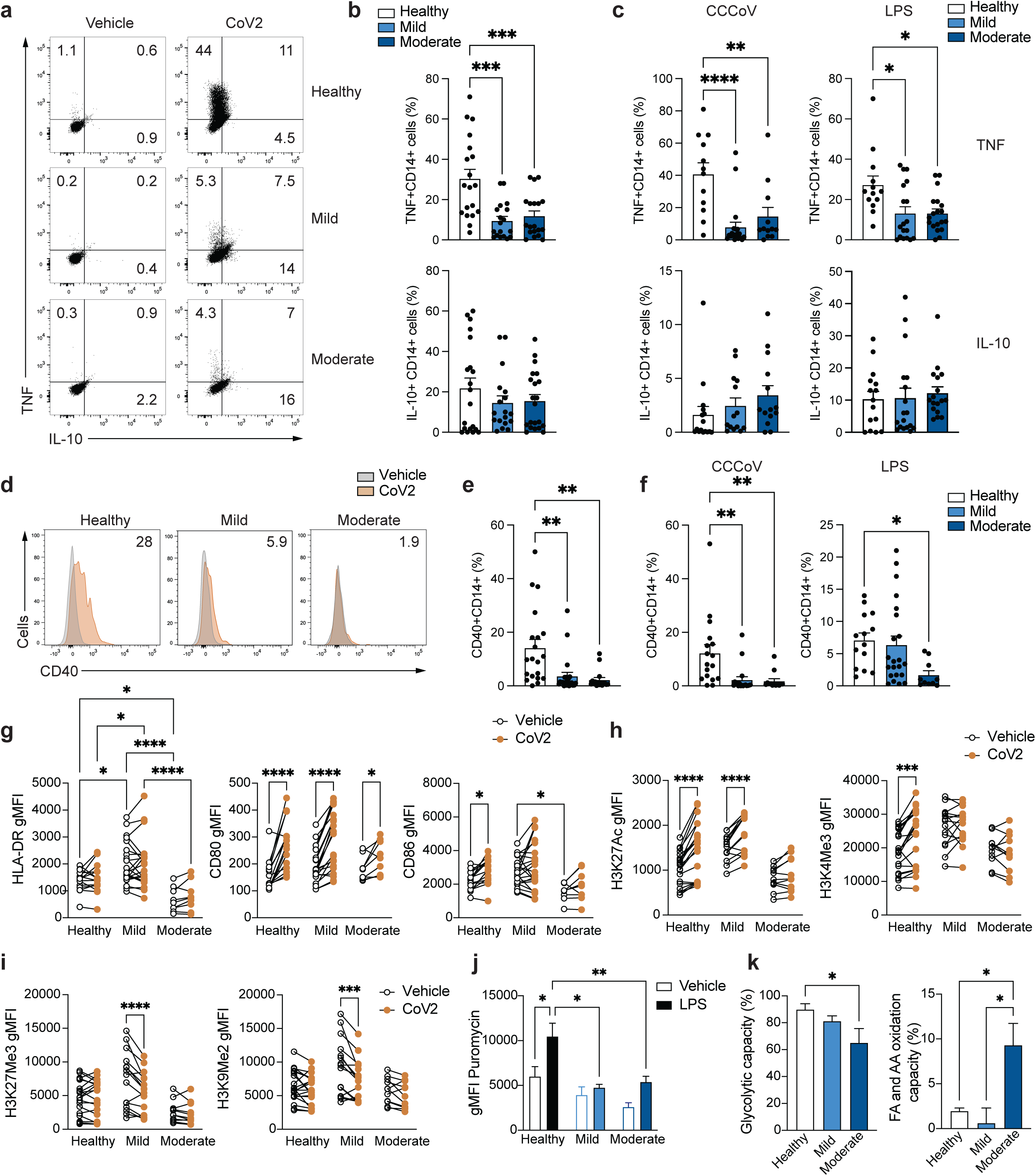
Impaired *ex vivo* pathogen sensing by COVID-19 monocytes. **a**. Representative example of the production of TNF and IL-10 by CD14^+^ monocytes from healthy individuals, mild and moderate COVID-19 patients after *ex vivo* stimulation with SARS-CoV-2. **b**. Summary of percentage of TNF- and IL-10-producing CD14^+^ from CD14^+^ monocytes after SARS-CoV-2 stimulation in healthy individuals (n=19), mild (n=18) and moderate (n=19) COVID-19 patients. **c**. Summary of percentage of TNF- and IL-10-producing CD14^+^ from CD14^+^ cells after stimulation with a mixture of heat-inactivated common cold coronaviruses (CCCoV, left) or LPS (right) in healthy individuals (n=12 for CCCoV and n=13 for LPS stimulation), mild (n=21 for CCCoV and n=18 for LPS stimulation) and moderate (n=12 for CCCoV and n=19 for LPS stimulation) COVID-19 patients. **d**. Representative histograms of CD40 expression by healthy individual, mild and moderate COVID-19 monocytes stimulated with vehicle (grey histogram) or SARS-CoV-2 (orange histogram). Numbers represent percentage of CD40^+^ monocytes relative to vehicle-stimulated cells. **e**. Summary of percentage of CD40^+^CD14^+^ from CD14^+^ cells after SARS-CoV-2 stimulation in healthy individuals (n=20), mild (n=22) and moderate (n=16) COVID-19 patients. **f**. Summary of percentage of CD40^+^CD14^+^ from CD14^+^ cells after stimulation with a mixture of heat-inactivated common cold coronaviruses (CCCoV, left) or LPS (right) in healthy individuals (n=17 for CCCoV and n=14 for LPS stimulation), mild (n=18 for CCCoV and n=22 for LPS stimulation) and moderate (n=13 for CCCoV and n=10 for LPS stimulation) COVID-19 patients. **g**. Summary of HLA-DR (left), CD80 (middle) and CD86 (right) expression measured by flow cytometry and plotted as gMFI of CD14^+^ monocytes from healthy individuals (n=15), mild (n=22) and moderate (n=9) COVID-19 patients stimulated with vehicle (white dots) or SARS-CoV-2 (CoV2, orange dots). Lines link paired samples. **h**. Summary of H3K27Ac (left) and H3K4Me3 (right) expression measured by flow cytometry and plotted as gMFI of CD14^+^ monocytes from healthy individuals (n=20), mild (n=15) and moderate (n=11) COVID-19 patients stimulated with vehicle (white dots) or SARS-CoV-2 (CoV2, orange dots). Lines link paired samples. **i**. Summary of H3K27Me3 (left) and H3K9Me2 (right) expression measured by flow cytometry and plotted as gMFI of CD14^+^ monocytes from healthy individuals (n=20), mild (n=15) and moderate (n=11) COVID-19 patients stimulated with vehicle (white dots) or SARS-CoV-2 (CoV2, orange dots). Lines link paired samples. **j**. Energetic status measured by puromycin expression (gMFI) of monocytes from healthy individuals (n=10), mild (n=8) or moderate (n=10) COVID-19 patients stimulated with vehicle (open bars) or LPS (colored bars). **k**. Glycolytic capacity (%, left) and fatty acid and amino acid oxidation capacity (%, right) of CD14^+^ monocytes from healthy individuals (n=10), mild (n=8) and moderate (n=10) COVID-19 patients stimulated with LPS. One-way ANOVA with Tukey’s correction for multiples comparisons in **b, c, e, f** and **k**. Two-way ANOVA with Tukey’s correction for multiple comparisons in **g, h, i, j**. *p<0.05, **p<0.005, ***p<0.001, ****p<0.0001.

Epigenetic reprogramming underlies innate immune cell activation upon pathogen sensing. In agreement with this, monocytes from healthy individuals significantly increased the expression of H3K27Ac and H3K4Me3, associated with activation of gene expression^26,27^, upon SARS-CoV-2 stimulation. In contrast, monocytes from moderate COVID-19 patients did not change the expression of these histone marks after SARS-CoV-2 sensing. Monocytes from mild COVID-19 patients demonstrated an intermediate pattern of expression, with significant upregulation of H3K27Ac but no change in H3K4Me3 upon SARS-CoV-2 stimulation (Figure 3h). Moreover, mild patient monocytes significantly decreased the expression of repressive H3K27Me3 and H3K9Me2 marks, while neither healthy or moderate COVID-19 monocytes did after stimulation with SARS-CoV-2 (Figure 3i).

The apparent unresponsiveness of COVID-19 monocytes to pathogen sensing was accompanied by altered metabolic reprogramming. Innate immune cells that sense pathogens increase the rate of glycolysis over mitochondrial oxidative phosphorylation to enable fast energy availability ^37-39^. However, COVID-19 monocyte energetic profile measured by SCENITH™ did not increase upon LPS stimulation, unlike that of healthy monocytes (Figure 3j). Moreover, moderate COVID-19 monocytes showed a decreased glycolytic capacity and an increase in fatty acid and amino acid oxidation capacity (Figure 3k) compared to healthy monocytes, that correlated with a slight but significant decrease in glucose dependency and an increase in mitochondrial dependency compared to monocytes from healthy individuals (Supplementary Figure 5). These data are in agreement with the enriched metabolic pathways from RNA-seq data (Figures 2c and 2h). Seahorse experiments confirmed the defect in glycolysis in stimulated monocytes from COVID-19 patients (Supplementary Figure 6). In summary, monocytes from COVID-19 patients display a profound defect in pathogen sensing *ex vivo* that is more evident in moderate than in mild patients and is characterized by an impairment in pro-inflammatory cytokine production and expression of activation-related receptors, epigenetic reprogramming and metabolic rewiring.

### SARS-CoV-2-stimulated monocytes from COVID-19 patients display a pro-thrombotic gene expression signature

To globally characterize the gene expression signature of activated monocytes in COVID-19, we performed RNA-seq on SARS-CoV-2-stimulated monocytes from healthy individuals and patients with moderate COVID-19 (Figure 4). PCA clearly separated COVID-19 from healthy monocytes, although some healthy monocytes clustered with COVID-19 in the principal component space (Figure 4a, Supplementary Figure 7). Quantification of differentially expressed genes yielded 1,437 upregulated and 2,073 downregulated genes in activated COVID-19 compared to activated healthy monocytes (≥1.5 fold change, FDR<0.05, Figure 4b). Pathway enrichment of differentially expressed genes (≥1.5 fold change vs. healthy monocytes, FDR<0.05) using XGR software and the Reactome pathway database demonstrated a number of expected pathways involved in the innate immune response to pathogens, including type I IFN signaling, cytokine signaling, interactions between lymphoid and non-lymphoid cells, NLR sensing, etc (Supplementary Figure 8 and Supplementary Table 6). However, when we focused our analysis on pathways enriched in upregulated genes in activated COVID-19 monocytes compared to activated healthy monocytes, the most significantly enriched pathways were involved in hemostasis and coagulation, including integrin signaling, extracellular matrix organization, signaling by PDGF, interactions with activated platelets and general hemostasis (Figure 4c and Supplementary Table 7). Integrin receptors are used by cells to interact with other cells and with the extracellular matrix, by binding numerous matrix proteins including collagen, actin and laminin being also involved in hemostasis and platelet aggregation^40^. In addition, monocytes actively bind to platelets forming pro-thrombotic aggregates in inflammatory and vascular pathologies^41,42^. Monocytes from COVID-19 patients expressed increased levels of various collagen subunits (*COL1A1, PLOD2, COL6A3, COL6A1*), enzymes involved in collagen triple helix synthesis (*COLGALT1*) and a number of matrix metalloproteinases (*MMP1, MMP2, MMP14*, Figure 4d), which are not only involved in extracellular matrix remodeling, but they have also been implicated in contributing directly to platelet activation and priming for aggregation^43,44^. These results are in agreement with the clinical observations of hypercoagulability and acquired coagulopathies in patients with COVID-19^45-48^, and suggest that monocytes from moderate COVID-19 patients upregulate a pro-thrombotic gene expression signature upon further SARS-CoV-2 sensing.

**Figure 4.**
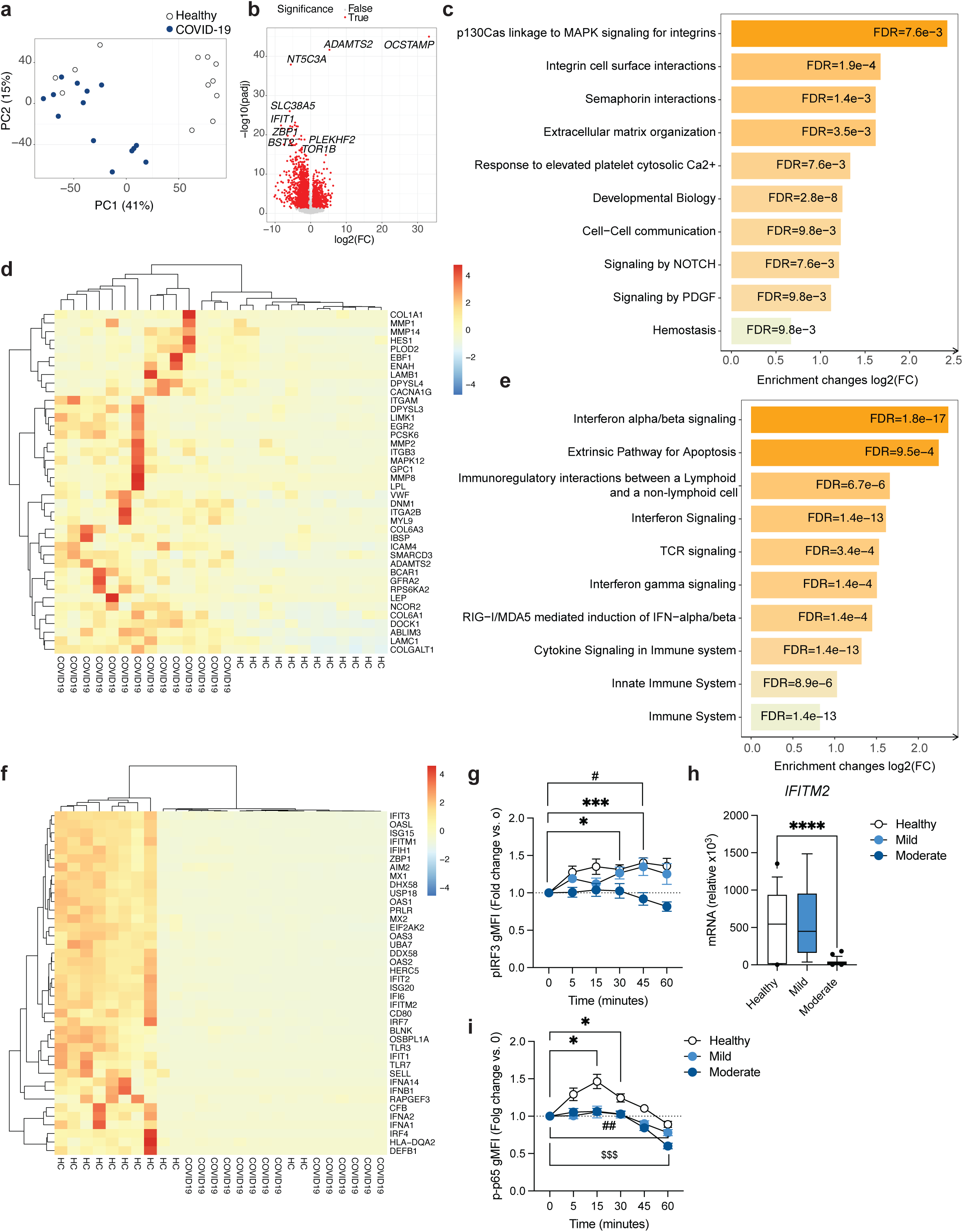
Gene expression signature of COVID-19 monocytes upon pathogen sensing. **a**. Principal component analysis (PCA) of the gene expression data computed from all genes from healthy individual (white dots) and moderate COVID-19 (blue dots) monocyte samples stimulated with SARS-CoV-2. PC2 plotted against PC1 to explore overall variation across samples. The variance explained by each component is stated in brackets. **b**. Volcano plots of differentially expressed genes for activated COVID-19 vs. activated healthy monocytes. Red coloring shows genes with fold change ≥ 1.5 and FDR<0.05. **c**. Bar plots depict the top 10 significantly enriched (FDR<0.05) pathways from Reactome for COVID-19 vs. healthy individual monocytes stimulated with SARS-CoV-2 using upregulated genes in COVID-19 vs healthy (≥ 1.5 fold increase, FDR<0.05), with the fold enrichment plotted on the x axis as log_2_ (FC) and the bars labelled with the adjusted p value. **d**. Top 40 significantly upregulated genes in the COVID-19 vs healthy monocyte contrast that are members of the pathways in **c**, shown in a heatmap. Gene expression values are scaled by row, with red indicating relatively high expression and blue low expression. Both rows and columns are clustered using Euclidean distance and Ward’s method. **e**. Bar plots depict the top 10 significantly enriched (FDR<0.05) pathways from Reactome for COVID-19 vs. healthy individual SARS-CoV-2-stimulated monocytes, using downregulated genes in COVID-19 vs healthy (≥ 1.5 fold decrease, FDR<0.05), with the fold enrichment plotted on the x axis as log_2_ (FC) and the bars labelled with the adjusted p value. **f**. Top 40 significantly downregulated genes in the SARS-CoV-2-stimulated COVID-19 vs. healthy individual monocyte contrast that are members of the pathways in **e**, shown in a heatmap. Gene expression values are scaled by row, with red indicating relatively high expression and blue low expression. Both rows and columns are clustered using Euclidean distance and Ward’s method. **g**. Phospho-IRF3 (Ser 396) expression measured by flow cytometry and plotted as fold change to baseline (gMFI) for healthy (n=14, white dots), mild (n=15, light blue dots) and moderate (n=10, dark blue dots) COVID-19 monocytes stimulated with LPS for 60 minutes. **h**. *IFITM2* relative gene expression (to *GAPDH*) measured by real-time PCR in sorted CD14^+^ monocytes from healthy individuals (n=14), mild (n=7) and moderate (n=23) COVID-19 stimulated with SARS-CoV-2. **i**. Phospho-NFκB p65 (Ser 529) expression measured by flow cytometry and plotted as fold change to baseline (gMFI) for healthy (n=14, white dots), mild (n=15, light blue dots) and moderate (n=10, dark blue dots) COVID-19 monocytes stimulated with LPS for 60 minutes. Mixed model with Tukey’s post-test for multiple comparisons for **g** and **i**. One-way ANOVA with Tukey’s test for multiple comparisons in **h**. For **g** and **i**, statistical significance of only baseline vs. other time points within the same patient groups are shown. *p<0.05, ***p<0.001 for healthy individual comparisons, #p<0.05, ##p<0.005 for mild COVID-19 patient comparisons, $$$p<0.001 for moderate COVID-19 patient comparisons. ****p<0.0001.

Interestingly, downregulated pathways in stimulated COVID-19 monocytes included most of the canonical immunological functions expected for innate immune cells upon virus sensing, i.e. interferon signaling, RIG-I/MDA5-mediated induction of interferons, activation of TCR signaling in T cells, innate immune functions and interactions with non-lymphoid cells (Figure 4e and Supplementary Table 8). The majority of the top 40 genes significantly downregulated in COVID-19 monocytes from these downregulated pathways consisted of different interferons (*IFNA1, IFNA2, IFNA14* and *IFNB1*), interferon-stimulated genes (*IFIT3, ISG15, IFIT2, ISG20, IRF7* and *MX2*) and pathogen-sensing receptors (*TLR7, AIM2*, Figure 4f). This gene signature was functionally confirmed by examining the activation pattern of IRF3 in response to LPS in monocytes from healthy individuals and patients with mild and moderate COVID-19 (Figure 4g). While healthy and mild COVID-19 monocytes significantly increased the expression of the phosphorylated form of IRF3 upon LPS stimulation compared to baseline levels, monocytes from moderate patients did not. This inability to activate IRF3 correlated with decreased expression of the interferon-stimulated gene *IFITM2*, examined in an expanded cohort of healthy, mild and moderate COVID-19 monocytes after stimulation with SARS-CoV-2 (Figure 4h). Of note, examination of NFκB p65 activation, as a main transcription factor involved in cytokine signaling in innate cells, demonstrated a defective activation in both mild and moderate COVID-19 as compared to healthy individuals (Figure 4i).

These findings are consistent with an unexpected transcriptional and functional switch of COVID-19 monocytes from canonical innate immune functions to a pro-thrombotic phenotype and potential cross-talk with other cells involved in hemostasis, which suggests that activated monocytes may contribute to COVID-19 severity by actively impacting hemostasis and by a reduction in innate immune functions necessary for efficient virus clearance.

### Endotoxin tolerance signature enriched in activated COVID-19 monocytes

A number of works have suggested similarities between the characteristics of the immune response in COVID-19 patients and those of septic individuals, including multiple organ dysfunction, immunosuppression, coagulopathies and acute respiratory failure^49^. To determine the similarities between the transcriptional signature of COVID-19 monocytes with that of sepsis monocytes, we utilized publicly available microarray gene expression data on sepsis monocytes and healthy controls^50^ and we tested the estimated fold changes for correlation with those from our *ex vivo* (Figure 5a) and activated (Figure 5b) COVID-19 and healthy monocytes. No clear correlation was observed in any of the two contrasts, which suggest that the transcriptional signature of CD14^+^ monocytes in moderate COVID-19 is not similar to that of monocytes in sepsis.

**Figure 5.**
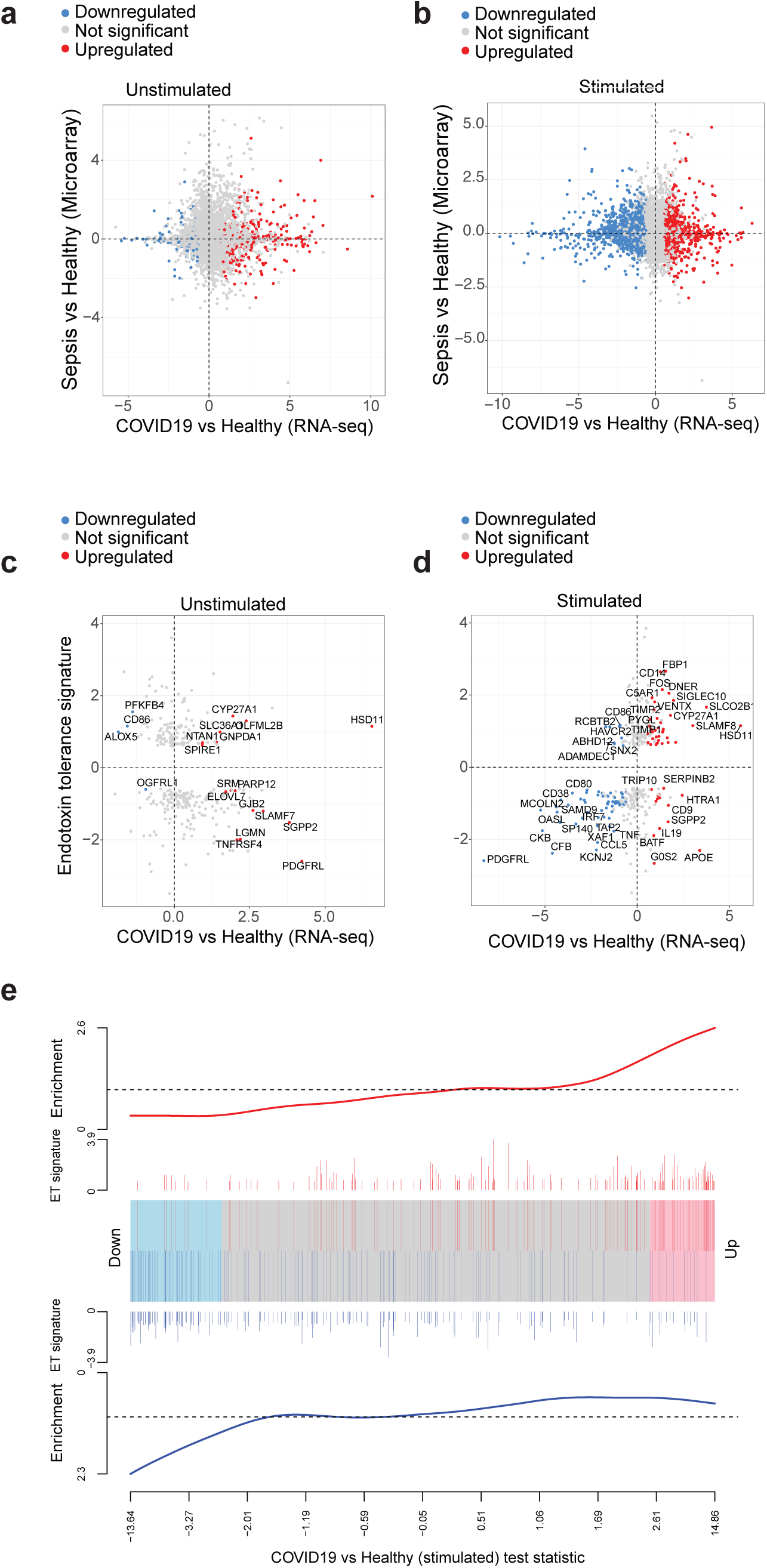
Endotoxin-induced tolerance signature significantly enriched in COVID-19 monocytes. **a**. Correlation plot of sepsis vs. healthy individual gene expression signature and *ex vivo* COVID-19 vs. healthy individual monocyte gene expression signature. Each point represents a gene detected in both the public sepsis dataset and our COVID-19 RNA-seq dataset. The log_2_FC between sepsis and healthy controls is plotted against the log_2_FC for *ex vivo* COVID-19 monocytes vs. healthy control monocytes, and the points are colored according to the significance and direction of effect in the COVID-19 contrast (grey, not significant; red, significantly upregulated, blue, significantly downregulated). **b**. Correlation plot of sepsis vs. healthy individual gene expression signature and SARS-CoV-2-stimulated COVID-19 vs. healthy individual monocyte gene expression signature. **c**. Correlation plot of endotoxin-induced tolerance gene signature and *ex vivo* COVID-19 vs. healthy monocyte signature. Each point represents a gene detected in both the endotoxin gene signature and our COVID-19 vs. healthy RNA-seq dataset. The log_2_FC between endotoxin tolerance and LPS-response is plotted against the log_2_FC for *ex vivo* COVID-19 vs. healthy monocytes, and the points colored according to the significance and direction of effect in the COVID-19 contrast. Some of the most differentially expressed genes in the COVID-19 vs. healthy monocyte dataset are identified in the plot. **d**. Correlation plot of endotoxin-induced tolerance gene signature and SARS-CoV-2-stimulated COVID-19 vs. healthy monocyte signature. Each point represents a gene detected in both the endotoxin gene signature and our COVID-19 vs. healthy RNA-seq dataset. The log_2_FC between endotoxin tolerance and LPS-response is plotted against the log_2_FC for SARS-CoV-2-stimulated COVID-19 vs healthy monocytes, and the points colored according to the significance and direction of effect in the COVID-19 contrast. Some of the most differentially expressed genes in the COVID-19 vs. healthy monocyte dataset are identified in the plot. **e**. Barcode plot showing enrichment of the endotoxin tolerance gene set (ET) in the differential gene expression results for SARS-CoV-2-stimulated COVID-19 vs healthy monocytes. The ranked test statistics from DESeq2 for the SARS-CoV-2-stimulated COVID-19 vs. healthy contrast are represented by the central shaded bar, with genes downregulated in COVID-19 on the left and upregulated genes on the right. The ranks of the endotoxin tolerance gene set within the COVID-19 contrast are indicated by the vertical lines in the central bar. The weights of the endotoxin tolerance genes (log_2_ (FC) from the ET differential expression analysis) are indicated by the height of the red and blue lines above and below the central bar. The red and blue lines at the top and bottom indicate relative enrichment of the endotoxin tolerance genes (split into genes with positive and negative FCs in the ET contrast) in each part of the plot.

The lack of cytokine expression, activation of costimulatory receptors, impaired antigen presentation potential and metabolic impairments displayed by moderate COVID-19 monocytes resembled the phenotype observed in LPS-induced tolerance^51^. We have previously defined an endotoxin tolerance gene expression signature from publicly available microarray data on monocytes stimulated *in vitro* with LPS^52^ that comprises 398 genes. Out of these, 318 genes were detected in our RNA-seq dataset. We tested for correlation of the endotoxin tolerance signature with *ex vivo* (Figure 5c) and activated (Figure 5d) COVID-19 monocytes, and while *ex vivo* COVID-19 monocytes did not display a clear correlation with the tolerance signature, activated COVID-19 monocytes displayed similar directionality of expression in those genes from the tolerance signature that were detected in the dataset. These data were further confirmed in barcode plots (Figure 5e), showing a statistically significant enrichment of the endotoxin tolerance gene signature in the list of differentially expressed genes from stimulated COVID-19 monocytes compared to healthy controls, for both upregulated and downregulated genes.

## Discussion

Here we employed metabolic, transcriptomic and functional assays to identify a number of phenotypic and functional alterations of COVID-19 monocytes that characterize moderate disease and we have provided the functional characteristics of monocyte responses in mild SARS-CoV-2 infections as an example of an efficiently and successfully cleared infection without excessive immunopathology. Important alterations in epigenetic marks, metabolism and transcriptional signatures characterize moderate COVID-19 monocytes and are important aspects of a global unresponsiveness phenotype upon pathogen sensing characterized by a transcriptional switch from canonical innate immune functions to a pro-thrombotic signature. Epigenetic and metabolic defects probably underlie the observed dysfunctional phenotype as they modulate innate immune functions including cytokine expression, activation, phagocytic capacity, etc^34,53,54^. Moreover, it would be plausible that these two mechanisms are interlinked. For example, the defects in histone acetylation could be due to a lack of acetyl groups, which are mostly provided by acetyl-CoA generated as a glycolysis product^55^, which is inhibited in COVID-19 monocytes (Figures 2 and 3).

A question that remains to be answered is the driver(s) of the described circulating monocyte dysfunction. *Ex vivo*, pathogen sensing triggers a switch in COVID-19 monocyte gene expression signature from canonical innate immune functions to pro-thrombotic phenotype. It remains to be determined whether other soluble factors in the microenvironment contribute to this reprogramming, or even the direct infection of monocytes by SARS-CoV-2, which has been previously suggested^56^. The phenotype we observed in circulating monocytes is in clear contrast with the functionality of monocyte-derived macrophages in the lung of COVID19 patients^10^. In this regard, our study is limited by the lack of bronchoalveolar lavage fluid (BALF) paired samples to compare the phenotype and function of circulating monocytes with those infiltrating the target tissue. However, some previous publications examining paired airway and blood samples have shown differences in the signatures of circulating and lung innate immune cells, with low HLA-DR expressing, dysfunctional monocytes in the blood and hyperactive airway monocyte and macrophages producing pro-inflammatory cytokines^10,33,57^. The underlying mechanisms for these differences remain elusive. During the course of viral infections, circulating monocytes rapidly leave the bloodstream and migrate to target tissues, where after pathogen sensing and/or other microenvironmental stimuli, they differentiate into macrophages and/or dendritic cells. In this study we examined the functionality of monocytes during the acute phase of disease, early after symptom onset. It remains to be determined whether these dysfunctional monocytes have the capacity to migrate to the lungs and contribute to lung inflammation, or whether their dysfunction is such that migration is impaired and monocyte migration only occurred during the very initial phases of infection before monocyte acquired the impairments observed in this study. Of note, some of the defective pathways displayed by COVID-19 monocytes, as for example glycolysis, have been shown to be essential for migration of other cells to target tissue^58,59^. Finally, the results described in this study beg the question of whether the functional impairments observed in monocytes during the acute phase of infection are COVID-19-specific. While stimulation with other viruses and bacterial products led to similar altered immune phenotypes in COVID-19 monocytes (Figure 3), it seems likely that these processes occur with other moderate respiratory viral infections, as is the case during seasonal Influenza vaccination^60^. Longitudinal studies of monocyte dynamics during SARS-CoV-2 and other respiratory viral infections using both blood and BALF samples are warranted to answer these questions.

## Tables

Supplementary Table 1. Participant characteristics.

Supplementary Table 2. Percentage of cells per cluster in each study group.

Supplementary Table 3. Pathway enrichment of all differentially expressed genes from COVID-19 vs. healthy monocytes.

Supplementary Table 4. Pathway enrichment of upregulated genes from COVID-19 vs. healthy monocytes.

Supplementary Table 5. Pathway enrichment of downregulated genes from COVID-19 vs. healthy monocytes.

Supplementary Table 6. Pathway enrichment of all differentially expressed genes from stimulated COVID-19 vs. stimulated healthy monocytes.

Supplementary Table 7. Pathway enrichment of upregulated genes from stimulated COVID-19 vs. stimulated healthy monocytes.

Supplementary Table 8. Pathway enrichment of downregulated genes from stimulated COVID-19 vs. stimulated healthy monocytes.

## Figure legends

**Supplementary Figure 1.**
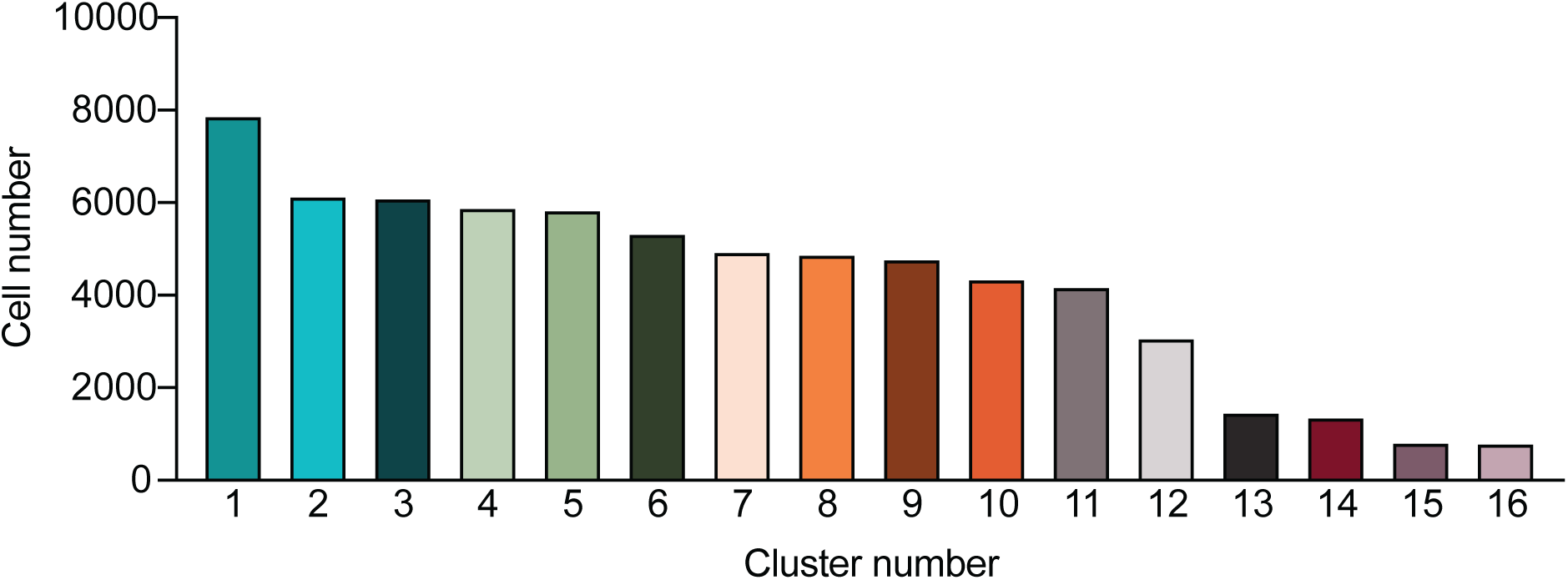
Number of cells per cluster identified by Phenograph.

**Supplementary Figure 2.**
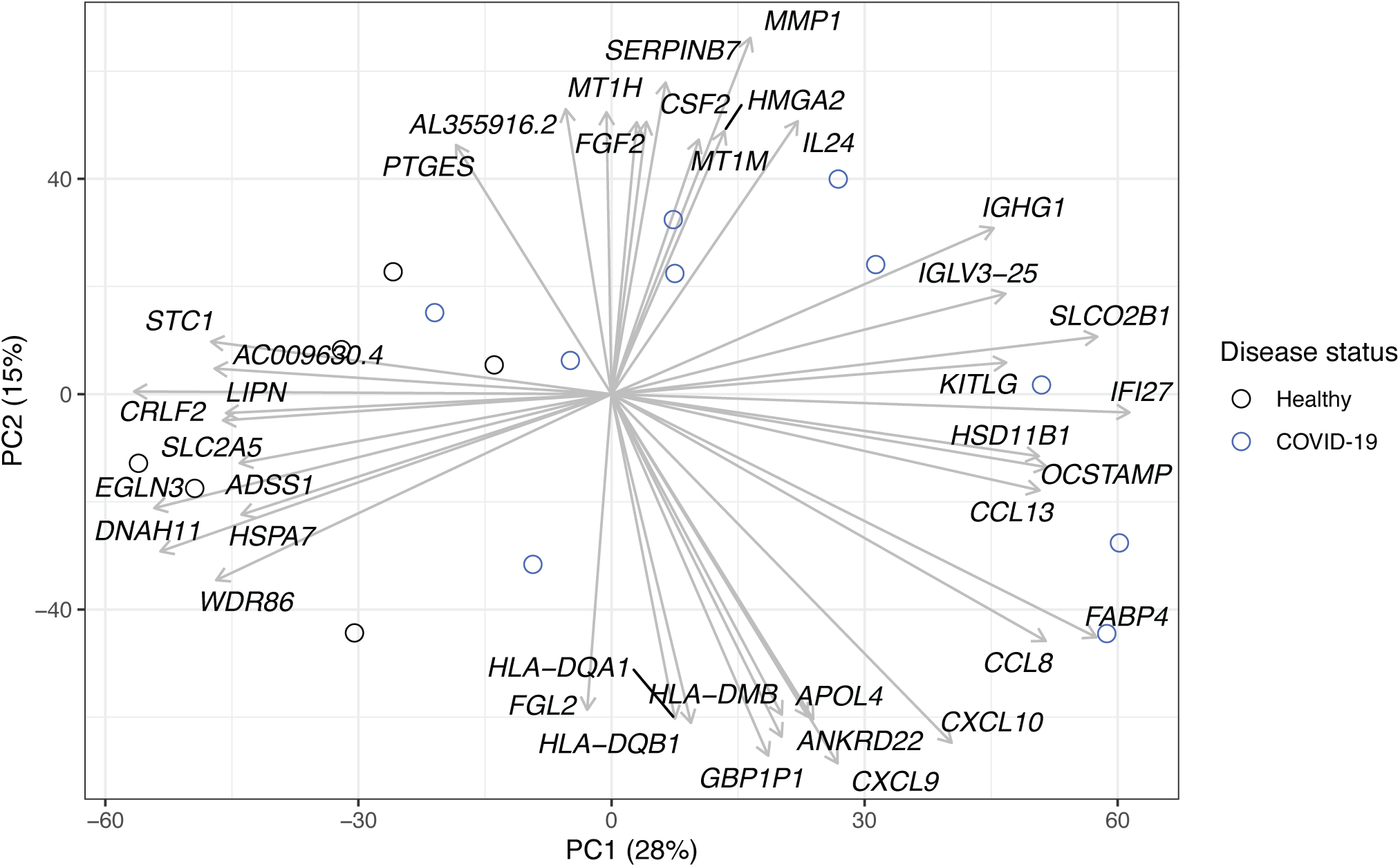
PCA gene loadings for RNA-seq of *ex vivo* isolated CD14^+^ monocytes from healthy individuals and moderate COVID-19 patients. The features contributing most to PC1 and PC2 (both positively and negatively) were identified using gene loadings, and the top 10 features for each PC are indicated, with arrows drawn from the origin illustrating their relative weights.

**Supplementary Figure 3.**
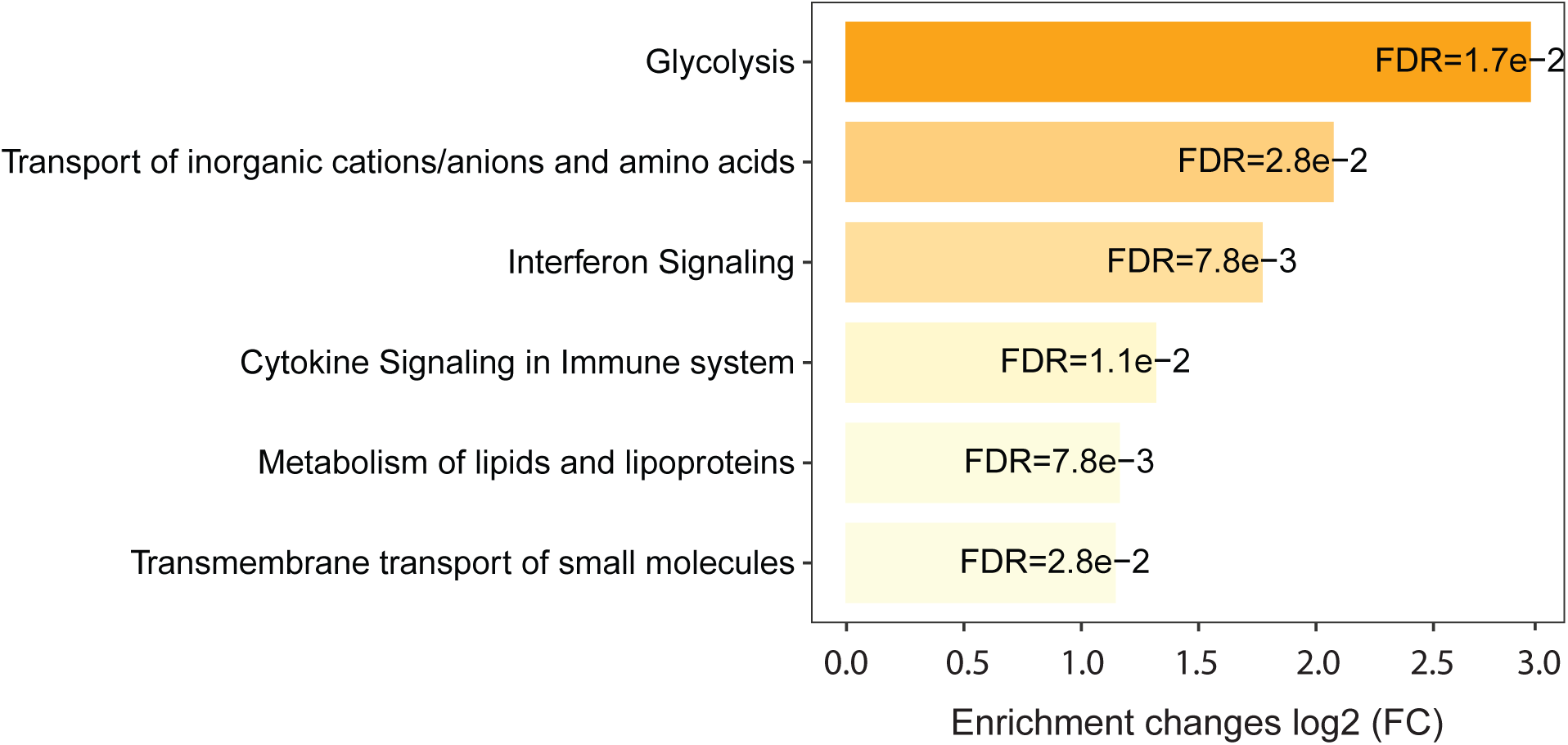
Pathway enrichment of COVID-19 monocyte RNA-seq data. Significantly enriched (FDR <0.05) pathways from Reactome for the *ex vivo* COVID-19 *vs*. healthy control monocytes differentially expressed genes are displayed as a bar plot, with the fold enrichment plotted on the x axis (log2(FC)) and the bars labelled with the adjusted p value.

**Supplementary Figure 4.**
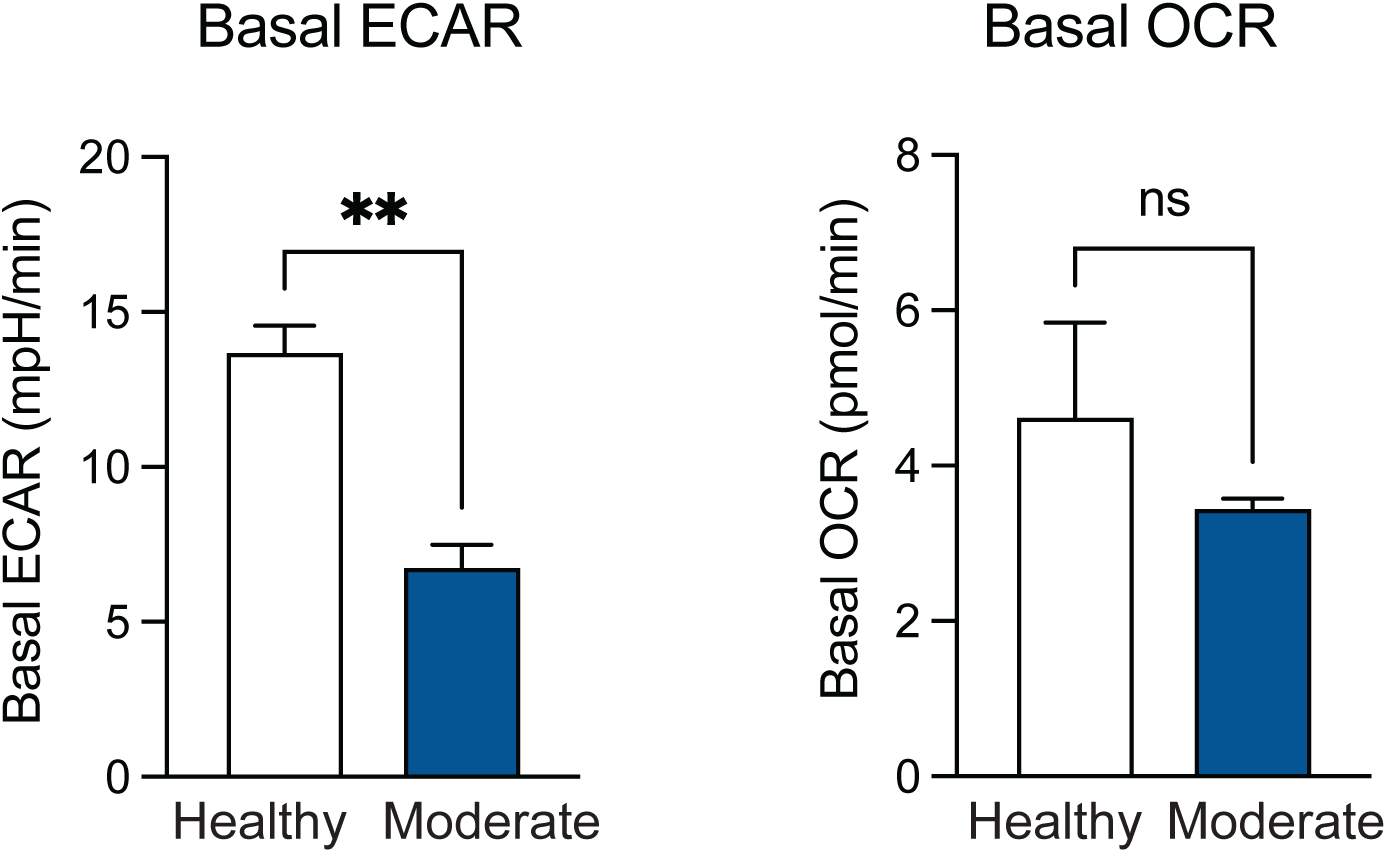
Seahorse analysis of COVID-19 monocytes *ex vivo*. Basal extracellular acidification rate (ECAR, left) and basal oxygen consumption rate (OCR, right) were measured in sorted CD14+ monocytes from healthy individuals (n=5) and COVID-19 patients (n=5). **p<0.005 by paired t-test.

**Supplementary Figure 5.**
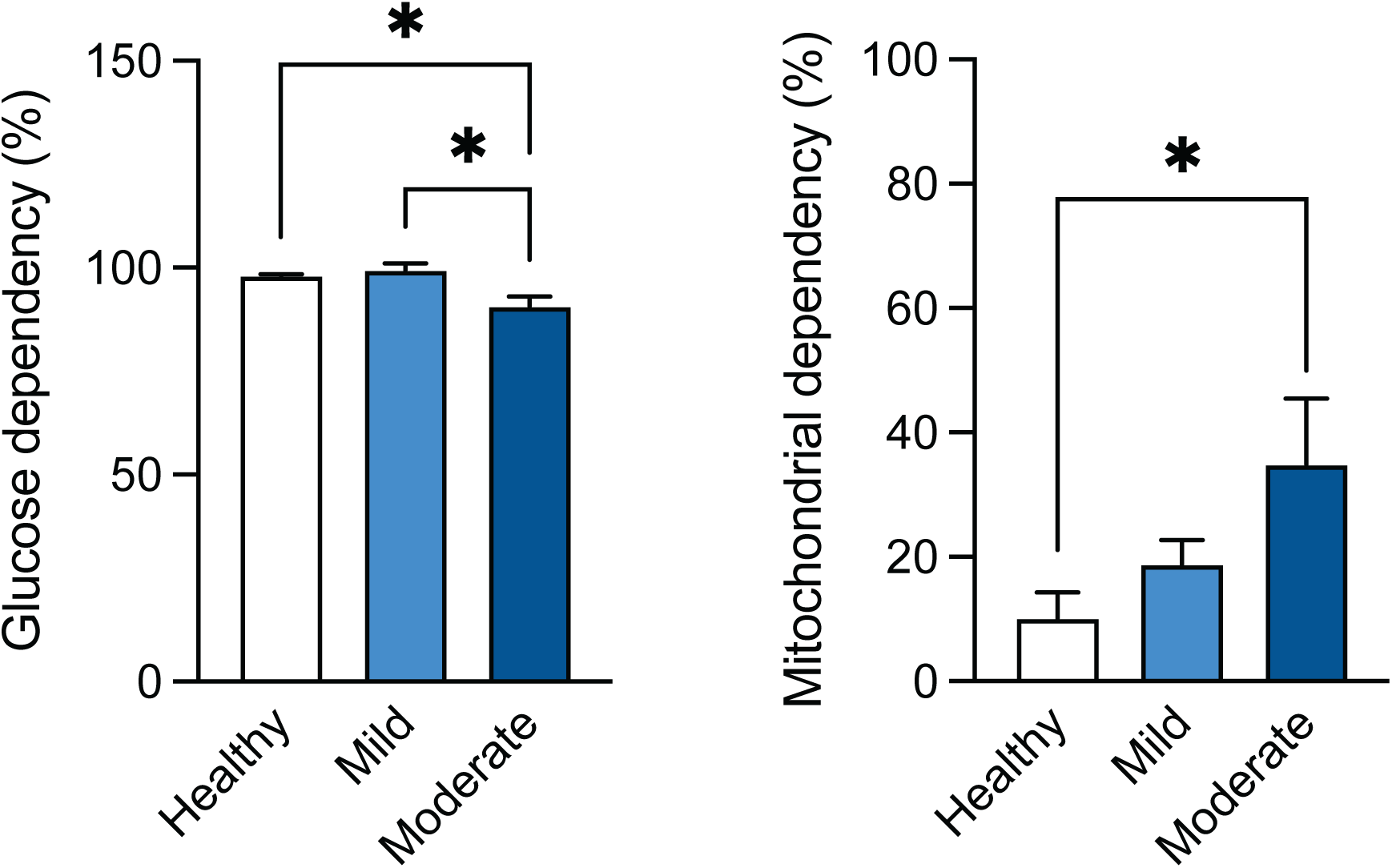
*Ex vivo* monocyte glucose metabolism and mitochondrial oxidation dependency. Glucose dependency (left) and mitochondrial oxidation dependency (right) calculated using SCENITH™ in healthy individuals (n=10, white bar), mild (n=8, light blue bar) and moderate (n=10, dark blue bar) COVID-19 monocytes.

**Supplementary Figure 6.**
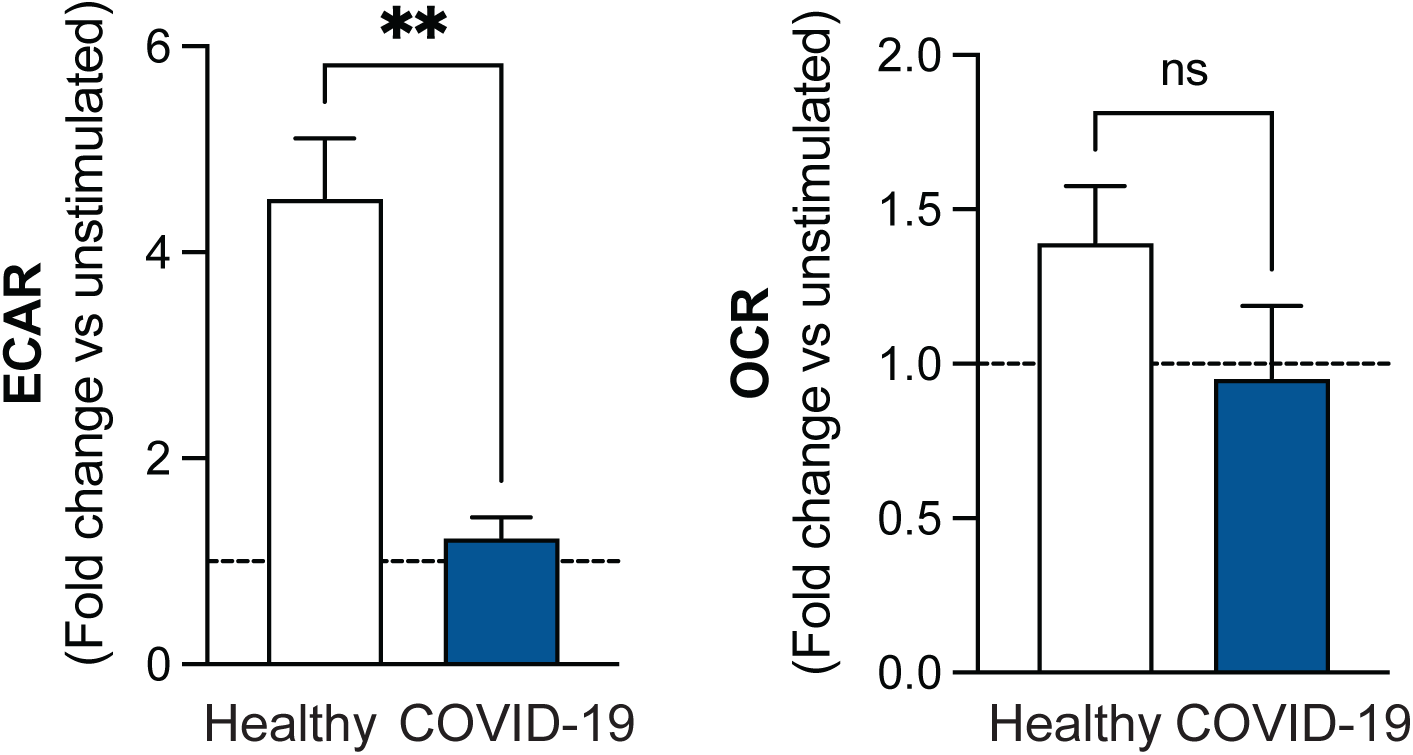
Seahorse analysis of activated COVID-19 monocytes. Extracellular acidification rate (ECAR, left) and oxygen consumption rate (OCR, right) were measured in sorted CD14^+^ monocytes from healthy individuals (n=5) and COVID-19 patients (n=5) stimulated or not with 100 ng/ml LPS for 18 hours. ECAR and OCR shown as fold increase relative to unstimulated controls **p<0.005 by paired t-test.

**Supplementary Figure 7.**
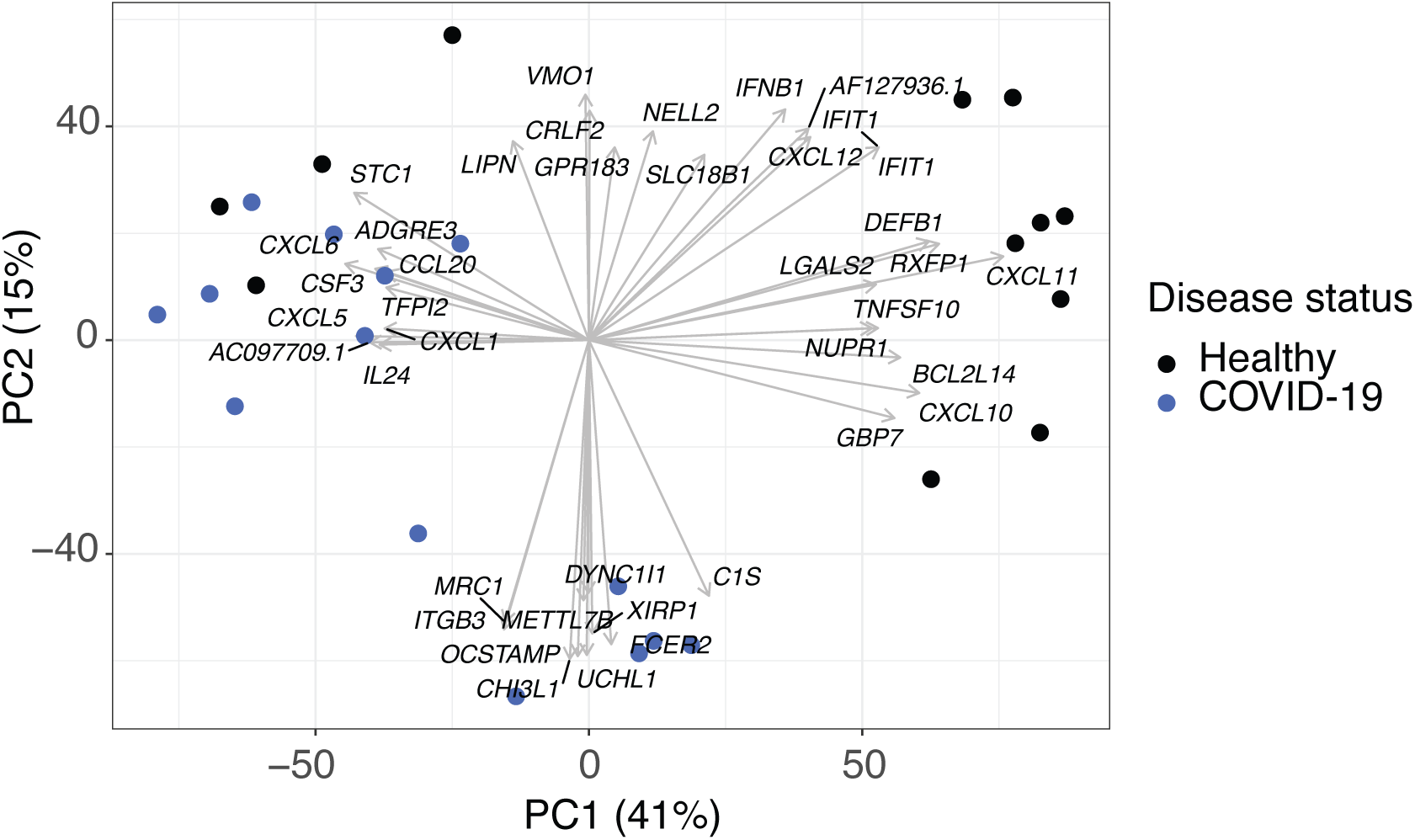
PCA gene loadings for RNA-seq of SARS-CoV-2-stimulated CD14^+^ monocytes from healthy individuals and moderate COVID-19 patients. The features contributing most to PC1 and PC2 (both positively and negatively) were identified using gene loadings, and the top 10 features for each PC are indicated, with arrows drawn from the origin illustrating their relative weights.

**Supplementary Figure 8.**
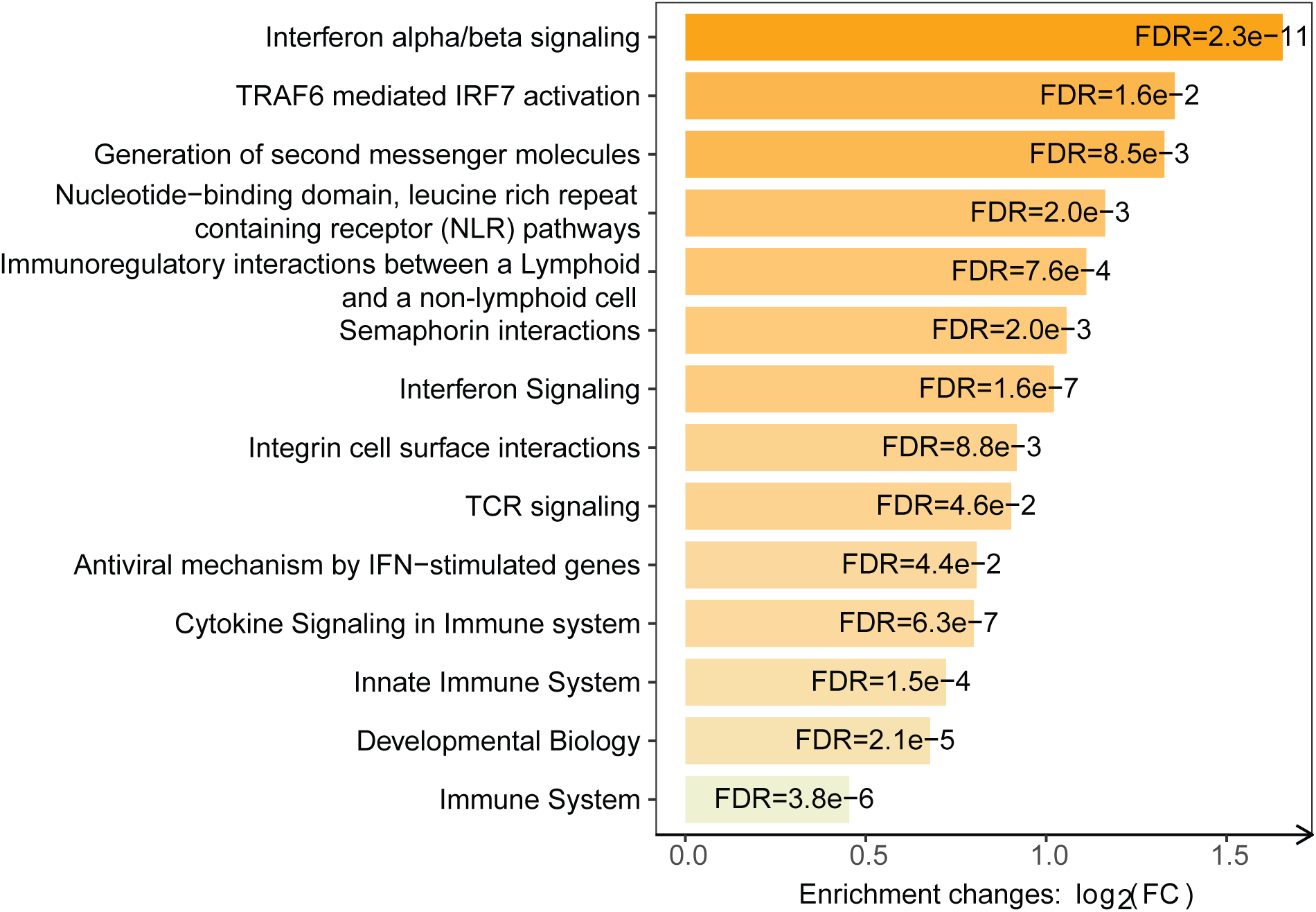
Pathway enrichment of SARS-CoV-2-stimulated COVID-19 monocyte RNA-seq data. Significantly enriched (FDR <0.05) pathways from Reactome for SARS-CoV-2 COVID-19 *vs*. healthy control monocytes differentially expressed genes are displayed as a bar plot, with the fold enrichment plotted on the x axis (log2(FC)) and the bars labelled with the adjusted p value.

## Materials and Methods

### Participants and clinical data collection

Disease severity was categorized based on the WHO ordinal classification of clinical improvement, where 0 (uninfected) describes people with no clinical or virological evidence of infection, 1-2 describe ambulatory patients without (1) or with (2) limitation of activities, and 3-4 corresponds to hospitalized patients with no oxygen therapy (3) or oxygen by mask or nasal prongs (4). Peripheral blood was collected from all participants and processed following a common standard operating protocol. For inpatients, clinical data were abstracted from the electronic medical records into summary participant sheets. Participant group characteristics are summarized in Supplementary Table 1.

Healthy donors (WHO 0) were Imperial College staff with no prior diagnosis of or recent symptoms consistent with COVID-19, and where possible, were matched in age and sex distribution with COVID-19 patients.

Blood samples from the COVID-19 patients examined in this work come from two different studies. COVIDITY study is a prospective observational serial sampling study of whole blood to observe the evolution of SARS-CoV-2 infection to characterize the host response to infection over time in peripheral blood (ethics approval obtained from the Health Research Authority, South Central Oxford C Research Ethics Committee). The population of study were >18 year old patients and/or staff at Imperial College Healthcare NHS Trust/Imperial College London with confirmed COVID-19 from a positive SARS-CoV-2 RT-PCR testing from NHS laboratories or Public Health England. Samples were taken 3-14 days after symptom initiation and were classified as 1 or 2 disease severity.

Samples from patients with moderate COVID-19 admitted to hospitals in London (Hammersmith Hospital, Charing Cross Hospital, Saint Mary’s Hospital) and eligible to participate in the MATIS trial^61^ provided consent (ethics approval by the Health Research Authority, London-Surrey Borders Research Ethics Committee) and blood was collected 3-14 days after disease onset and 0-2 days after hospitalization and positive PCR, and before study treatment initiation. Moderate patients displayed mild of moderate COVID-19 pneumonia, defined as grade 3 or 4 WHO severity. Samples were collected from March 2020 to February 2021 and none of the participants had received a COVID-19 vaccine.

### Cell Isolation and storage

Peripheral blood mononuclear cells (PBMCs) were isolated by Ficoll Hypaque (GE Healthcare) gradient centrifugation <4 hours after blood collection. The PBMC layer was collected, washed with PBS, resuspended at 20 million cells/ml in fetal bovine serum supplemented with 10% DMSO and stored at −150 ºC or liquid nitrogen.

### Flow cytometry stainings

PBMCs were thawed and rested for 2 hours at 37 ºC in RPMI 1640 media supplemented with 2 mM L-glutamine, 5% human AB serum, and 1x Penicillin and Streptomycin. For *ex viv*o phenotypic characterization, 300,000-500,000 PBMC were stained with LIVE/DEAD Fixable Dead Cell Dyes (Thermo Fisher Scientific) according to the manufacturer’s specifications. A Fc receptor (FcR) blocking step was performed using FcR Blocking Reagent Human (Miltenyi Biotec) before cell surface antibody staining. The antibodies used in the stainings were the following: CD14 (61D3, eBioscience), CD3 (UCHT1, BD), CD19 (HIB19, BD), CD1c (L161, Biolegend), CD40 (5C3, Biolegend), CD141 (M80, Biolegend), CD304 (12C2, Biolegend), CD86 (BU63, Biolegend), CD80 (BB1, BD Pharmigen), HLA-DR (L243, Biolegend), CD301 (H037G3, Biolegend), HLA-ABC (W6/32, Biolegend), TIM-3 (F38-2E2, Invitrogen), PD-1 (EH12.2H7, Biolegend), and CD16 (3G8, BD). Cells were subsequently fixed using the Foxp3 staining buffer kit (Thermo Fisher Scientific) following the manufacturer’s recommendations and resuspended in 250 ul of PBS.

For intracellular staining, the abovementioned protocol was used and an additional step for intracellular staining was added after fixation. The antibodies used for intracellular staining were the following: H3K27Ac, H3K9Me2, H3K4Me3, H3K27Me3 (all from Cell Signaling Technology), TNF (Mab11, Biolegen) and IL-10 (JES3-907, Thermo Fisher Scientific). Intracellular staining was performed using the the Foxp3 staining buffer kit.

Samples were run on a Fortessa instrument (BD Biosciences) and analyzed using FlowJo v.10. Dimensionality reduction and tSNE plots were obtained by downsampling each of the 15 samples per group (healthy, mild COVID-19 and moderate COVID-19) to 1,500 events per sample, and the concatenated sample was used to calculate tSNE axes using 1,000 iterations, perplexity of 40 and the default learning rate (4734). In order to obtain cell clusters, we used Phenograph^62^ plugin in FlowJo, with k=166 and all compensated parameters.

### Generation of virus stocks

SARS-CoV-2 virus (SARS-CoV-2/England/IC19/2020 isolate, kindly provided by Wendy S Barclay) was expanded in Vero-E6 cells. Briefly, Vero-E6 cells were plated in serum-free medium (OptiPRO SFM containing 2x GlutaMAX) in T75 flasks and infected with SARS-CoV-2 at a multiplicity of infection of 0.1 and a final volume of 5 ml. Cells were incubated for 2 hours at 37 °C, 5% CO_2_, after which the inoculum was removed and complete medium without serum was added to the culture. Cells were incubated for 3-5 days (until cytopathic effects were observed). Subsequently, cell culture supernatant was collected, centrifuged at 1000 x*g*, 4 ºC for 15 minutes and transferred to a new 50 ml tube for a second centrifugation at 1000 x*g*, 4 ºC for 15 minutes. Viral supernatant was collected, filtered through 0.45 µm and an aliquot was taken for titration. The rest of the supernatant was UV-inactivated and concentrated using Retro-X concentrator (Takara Bio), following manufacturer’s recommendations and published protocols^63,64^.

Human coronaviruses (CCCoV) 229E, OC43 and NL63 strains (Public Health England) were expanded in MRC-5 (kindly provided by Dr Rob White, Imperial College London), BSC-1 (Public Health England) and LLCMK2 (Public Health England), respectively. Briefly, cell lines were plated in serum-free medium (DMEM, 1x non-essential amino acids) in T75 flasks and infected with CCCoV (229E, OC43 or NL63) at a multiplicity of infection of 0.1 and a final volume of 5 ml. Cells were incubated for 2 hours at 37 ºC, 5% CO_2_, after which the inoculum was removed and medium without serum was added to the culture. Cells were incubated for 3-5 days (until cytopathic effects were observed). Subsequently, cell culture supernatant was collected, centrifuged at 1000 x*g*, 4 ºC for 15 minutes and transferred to a new 50 ml tube for a second centrifugation at 1000 x*g*, 4 ºC for 15 minutes. Viral supernatant was collected, filtered through 0.45 µm and an aliquot was taken for titration. The rest of the supernatant was heat-inactivated and concentrated using Retro-X concentrator (Takara Bio), following manufacturer’s recommendations and published protocols^63,64^.

### Titration of virus stocks

For SARS-CoV-2 titration, samples were serially diluted in OptiPRO SFM, 2X GlutaMAX (1:10) and added to Vero cell monolayers for 1 hour at 37 °C, 5% CO_2_. The inoculum was subsequently removed and cells were overlayed with DMEM containing 0.2% w/v bovine serum albumin, 0.16% w/v NaHCO_3_, 10 mM HEPES, 2 mM L-Gutamine, 1X P/S and 0.6% w/v agarose. Plates were incubated at 37 °C, 5% CO_2_ for 3 days. The overlay was then removed and monolayers were stained with crystal violet solution for 1 hour at room temperature. Plates were washed with water, dried and virus plaques were counted.

For CCCoV titration, viral supernatants were serially diluted in DMEM, non essential amino acids (1:10) and added to MRC-5 (229E strain), BSC-1 (OC43 strain) or LLCMK2 (NL63 strain) cell monolayers for 1 hour at 37 °C, 5% CO_2_. The inoculum was subsequently removed and cells were overlayed with DMEM medium for 4-5 days (until cytopathic effects were observed). An endpoint dilution assay was used to determine viral infectivity titers^63^.

### Ex vivo stimulation assays

PBMC were thawed and rested for 2 hours at 37 ºC in complete media. 250,000 PBMC were plated in polysterene plates (Corning) to prevent unspecific stimulation of monocytes by adherence to the plastic plate^65^. Cells were stimulated with vehicle, UV-inactivated SARS-CoV-2 (CoV-2), 100 ng/ml LPS or a mixture of heat-inactivated common cold coronaviruses consisting of the 229E, OC43 and NL63 strains (CCCoV) at 10^6^ viral particles per 10^6^ cells for 20 hours. For intracellular stainings, GolgiStop™ (BD Biosciences) was added to the cultures 10 hours after stimulation for a total of 10 hours.

### RNA isolation, RNA quality control, and sample preparation for RNA-seq analysis

Sorted CD14^+^ monocytes from total PBMC either *ex vivo* or after a 20 hour stimulation with 10^6^ UV-inactivated SARS-CoV-2 viral particles per 10^6^ cells were lysed with RLT Plus buffer (QIAGEN). RNA was isolated using the RNeasy Micro Plus Kit (QIAGEN) following the manufacturer’s guidelines in Appendix D of the QIAGEN RNeasy handbook. RNA quality was quantified using the Agilent RNA 6000 Pico Kit (Agilent Technologies) following the manufacturer’s guidelines. RNA samples were stored at −80 ºC until further processing.

### RNA-seq analysis

RNA-sequencing was performed by the Oxford Genomics Centre. PolyA-enriched strand-specific libraries were prepared using NEBNext Ultra II Directional RNA Library Prep Kits (Illumina). All samples were pooled together and 150bp PE reads were sequenced on a Novaseq system, resulting in a median read count of 28M per sample.

Raw data was processed using the Sanger Nextflow RNA-seq pipeline (https://github.com/wtsi-hgi/nextflow-pipelines). Briefly, reads were aligned to the reference genome (GRCh38.99) using STAR v2.7.3^66^ in the two-pass mode (ENCODE recommended parameters) and gene expression was quantified using featureCounts^67^. Mapping statistics and quality control metrics from FastQC and RNA-SeQC^68^ indicated high data quality for all samples with no outliers detected. RNA-seq data analysis was performed in R v4.1 in Rstudio Server. Features that did not have at least 10 reads in at least 6 samples (the size of the smallest biological subgroup) were filtered out using the genefilter package^69^, resulting in a processed data set on 16,328 features. Principal component analysis (PCA) with the prcomp function was used to explore the relationship between samples, after the filtered gene counts were transformed using a regularized log transformation from the DESeq2^70^ package.

Differential gene expression analysis was carried out using DESeq2, comparing unstimulated monocytes from COVID-19 patients (n=10) to unstimulated monocytes from healthy controls (HC) (n=6), and SARS-CoV-2-stimulated monocytes from COVID-19 patients (n=14) to stimulated monocytes from HC (n=12). Genes with FDR<0.05 and a fold change (FC)≥1.5 were deemed significantly differentially expressed. Pathway enrichment analysis was performed using Fisher’s exact test in XGR^29^ with annotations from Reactome, using all genes retained in the processed RNA-seq data as the background, and employing the xEnrichConciser options. An adjusted p-value (BH FDR) threshold of 0.05 was used to identify significantly enriched pathways. Pheatmap package was used to draw heatmaps illustrating variation in gene expression across samples.

For testing the enrichment of the sepsis signature in our datasets, publicly available microarray gene expression data on sepsis patients and healthy controls were accessed using GEOquery (GSE46955)^50^. Gene expression between patients and controls was compared using limma^71^, for both the unstimulated and stimulated conditions. Subsequently, the estimated fold changes were tested for correlation with those from the COVID-19 vs HC results. Where multiple probes were available for the same gene in the microarray dataset, the top ranked probe was selected for the comparison.

For comparison to the endotoxin-induced tolerance signature, we have previously defined an endotoxin tolerance gene signature^72^ from publicly available microarray data on *in vitro* LPS-stimulated monocytes. Briefly, two datasets (GSE15219^52^ and GSE22248^73^) were accessed through GEO. Genes that were differentially expressed following a single LPS treatment (LPS response genes), and that were also differentially expressed between singly- and doubly-stimulated cells were identified. This resulted in an endotoxin tolerance gene signature comprising 398 genes, of which 318 were detected in the RNA-seq dataset. We tested for enrichment of this gene set in the COVID-19 versus healthy contrasts using the geneSetTest function and barcodeplot functions from limma.

### Quantification of mRNA expression by RT-PCR

Isolated RNA was converted to complementary DNA by reverse transcription (RT) with random hexamers and Multiscribe RT (TaqMan Reverse Transcription Reagents; Thermo Fisher Scientific). For *IFITM2* expression assays, the Hs00829485_sH probe was used from Thermo Fisher Scientific. The reactions were set up using the manufacturer’s guidelines and run on a StepOnePlue Real-Time PCR Machine (Thermo Fisher Scientific). Values are represented as the difference in cycle threshold (Ct) values normalised to *GAPDH* expression (Hs02786624_g1) for each sample as per the following formula: Relative RNA expression = (2-ΔCt) x 1000^74^.

### Metabolic profiling using SCENITH^™^

SCENITH^™^ is a flow cytometry-based method for profiling energy metabolism with single cell resolution^35^ *ex vivo* or after *in vitro* stimulation in sorted cells or complex cell mixtures. It uses puromycin incorporation to nascent proteins as a measurement for protein translation, which is tightly coupled to ATP production and therefore can be used as a readout for the energetic status of the cells at a given time.

PBMC were plated at 250,000-300,000 cells per well in 96 well plates and rested for 2 hours at 37 °C, 5% CO_2_ for *ex vivo* stainings, or rested for 2 hours and stimulated for 20 hours with 100 ng/ml LPS. Subsequently, cells were treated for 45 minutes at 37 ºC, 5% CO_2_ with Control (vehicle, Co), 100 mM 2-deoxy-D-glucose (DG, Sigma-Aldrich), 1 µM oligomycin (O, Sigma-Aldrich) or a combination of both drugs (DGO). 10 µg/ml puromycin was added to all conditions for the same amount of time. Cells were subsequently washed with room temperature PBS and stained for viability, cell surface markers and fixed as described above. Intracellular staining of puromycin was performed using the anti-puromycin monoclonal antibody (1:600 dilution, clone R4743L-E8) for 45 minutes at 4 ºC. The anti-puromycin antibody and metabolic inhibitors for SCENITH^™^ were kindly provided by Dr Argüello.

For the analysis of the energetic status of cells, puromycin geometric mean fluorescence intensity was analyzed in each of the four abovementioned conditions (Co, DG, O, DGO). To calculate the percentage of glucose dependence, the following formula was used: 100*((Co-DG)/(Co-DGO). Mitochondrial dependence (%) was calculated as 100*((Co-O)/(Co-DGO). Glycolytic capacity (%) was calculated as 100-Mitochondrial dependence. Fatty acid and amino acid oxidation capacity (%) was calculated as 100-Glucose dependence.

### Metabolic profiling using Seahorse

Sorted CD14^+^ monocytes from unstimulated or SARS-CoV-2-stimulated (20 hours at 37 ºC, 5% CO_2_) PBMC were plated at a range of 80,000-120,000 in duplicates for healthy and COVID-19 sample pairs, based on the minimum cell number obtained for each pair of samples in individual experiments. An XFp real-time ATP rate assay kit (Agilent Technologies) was used following manufacturer’s recommendations and samples were run in a Seahorse XF HS Mini Analyzer (Agilent Technologies). For basal oxygen consumption rate (OCR) and extracellular acidification rate (ECAR) measurements, 10 cycles were run and their average was taken as basal values per subject tested.

### Phosphorylation assays by flow cytometry

For *ex vivo* phosphorylation assays, thawed PBMC were plated at 250,000 cells per well in 96 well polypropylene plates and rested for 2 hours at 37 ºC, 5% CO_2_. PBMC were fixed with pre-warmed (37 ºC) Cytofix (BD Biosciences) for 20 minutes at 37 ºC, 5% CO_2_ and permeabilized with Perm III buffer (BD Biosciences) overnight at −20 ºC. Cultures were subsequently stained with CD3 (UCHT1, BD Biosciences), CD20 (H1, BD Biosciences), CD14 (M5E2, Biolegend), CD16 (B73.1, BD Biosciences), phospho-IRF3 (Ser 396, Bioss), phospho-NFkB p65 (Ser 529, BD Biosciences) in PBS for 1 hour at room temperature, washed with PBS and resuspended in 250 µl PBS.

For phosphorylation assays after LPS stimulation, PBMC were plated as above and stimulated with 100 ng/ml LPS for a total of 1 hour. Samples were fixed at 0, 5, 15, 30, 45 and 60 minutes after LPS addition for 20 min at 37 ºC, 5% CO_2_ and stained as above.

## Acknowledgements

We thank the participants who volunteered for this study and the clinical teams of the COVIDITY and MATIS studies for patient recruitment and blood collection. We thank Dr Parisa Amjadi and Ms Radhika Patel for their help with flow cytometry sorting. AKM is a Wellcome Trust PhD scholar. KLB and EED are funded by the Wellcome Trust [108413/A/15/D]. For the purpose of Open Access, the author has applied a CC BY public copyright license to any Author Accepted Manuscript version arising from this submission. We thank the Wellcome Sanger Institute’s Human Genetics Informatics (HGI) team for mapping the RNA-sequencing reads. This work was funded by a Rosetrees Trust grant to MDV (M971).

## Author contributions

AKM performed experiments, analyzed data and wrote the manuscript, KLB analyzed the RNA-seq data and wrote the manuscript, EJ performed experiments, LB prepared SARS-CoV-2 virus stocks, CS and NG performed experiments, CES and RQ provided patient samples, RA provided the SCENITH^™^ kit reagents and advised on SCENITH^™^ data analysis and interpretation, WSB provided SARS-CoV-2 virus stock, NC provided patient samples and advised on the clinical aspects of COVID-19, GPT provided COVID-19 patient samples and advised on the clinical aspects of COVID-19, EED supervised RNA-seq data analysis and wrote the manuscript, MDV designed the study, performed experiments, analyzed data, wrote the manuscript and obtained funding. All authors revised and contributed to the editing of the manuscript.

## Data availability

RNA-seq data will be available at the European Genome-Phenome Archive (EGA) upon manuscript acceptance.

